# Spatiotemporal dynamics of primary and motile cilia throughout lung development

**DOI:** 10.1101/2024.10.25.620342

**Authors:** Stephen Spurgin, Ange Michelle Nguimtsop, Fatima N. Chaudhry, Sylvia N. Michki, Jocelynda Salvador, M. Luisa Iruela-Arispe, Jarod A. Zepp, Saikat Mukhopadhyay, Ondine Cleaver

**Affiliations:** Department of Molecular Biology, University of Texas Southwestern Medical Center, 5323 Harry Hines Blvd., Dallas, Texas, USA 75390; Department of Pediatrics, University of Texas Southwestern Medical Center, 5323 Harry Hines Blvd., Dallas, Texas, USA 75390; Division of Pulmonary and Sleep Medicine, Department of Pediatrics, Children’s Hospital of Philadelphia, Philadelphia, Pennsylvania USA 19104; Department of Cell Biology, University of Texas Southwestern Medical Center, 6000 Harry Hines Blvd., Dallas, Texas, USA 75390; Department of Cell and Developmental Biology, Northwestern Feinberg School of Medicine, Chicago, Illinois, USA 60611

## Abstract

Cilia are specialized structures found on a variety of mammalian cells, with variable roles in the transduction of mechanical and biological signals (by primary cilia, PC), as well as the generation of fluid flow (by motile cilia). Their critical role in the establishment of a left-right axis in early development is well described, as is the innate immune function of multiciliated upper airway epithelium. By contrast, the dynamics of ciliary status during organogenesis and postnatal development is largely unknown. In this study, we define the progression of ciliary status within the endothelium, epithelium, and mesenchyme of the lung. Remarkably, we find that endothelial cells (ECs) lack PC at all stages of development, except in low numbers in the most proximal portions of the pulmonary arteries. In the lung epithelium, a proximodistal ciliary gradient is established over time, as the uniformly mono-ciliated epithelium transitions into proximal, multiciliated cells, and the distal alveolar epithelium loses its cilia. Mesenchymal cells, interestingly, are uniformly ciliated in early development, but with restriction to PDGFRα+ fibroblasts in the adult alveoli. This dynamic process in multiple cellular populations both challenges prior assertions that PC are found on all cells, and highlights a need to understand their spatiotemporal functions.

**Highlights:**

- Primary cilia are found broadly throughout early embryonic tissues.
- Primary cilia are observed in both epithelial and mesenchymal cells in the early lung.
- Pulmonary endothelial cells largely do not possess primary cilia during embryonic development.
- Differential multiciliation and loss of epithelial cilia in a proximal-distal axis.
- Maintenance of cilia in adult pulmonary PDGFRα+ fibroblasts.

## Introduction

Cilia are highly specialized, hair-like organelles found on a variety of mammalian cells. Cilia can be broadly defined as either *non-motile* primary cilia (PC), which facilitate signal transduction by sensing chemical, biological, or mechanical stimuli, or as *motile cilia,* which function to generate directional fluid flow.^1^ Whether or not a cell constructs a cilium—and which kind it builds—likely reflects a need for a specific ciliary function.

The essential function of cilia is evident from the very beginning of embryonic development. A foundational example is found at the embryonic node in mouse, where mechanisms of planar cell polarity ensure that motile cilia are aligned to shuttle the morphogen Nodal asymmetrically, dictating the left-right asymmetry of the body plan and ensuring the heart and pancreas are on the left side of the body, while the liver and gallbladder on the right.^2^ Beyond the establishment of overall internal organ sidedness, however, primary cilia have also been shown to be critical to the later development of several tissues, including heart,^3,4^ bone,^5–7^ kidney^8^ and brain.^9,10^

These later, organ-specific developmental roles are evidenced by the wide-ranging clinical presentations of the *ciliopathies* (diseases caused by mutations in known ciliary proteins), which cannot be fully explained by defective function in the primary cilia in the ventral node. From study of polycystic kidney disease (PKD), Bardet-Biedl syndrome, Meckel-Gruber syndrome, or Joubert’s syndrome, defects in ciliary proteins have been linked to a wide range of multisystem birth defects. These include neural tube defects, congenital heart defects, respiratory problems, polydactyly, and more.^11^ Building from clinical observation, increasingly strong pre-clinical evidence suggests that congenital heart disease is, at its core, a multigenic ciliopathy.^5–7^ In the developing neural tube, PC function as a sensory organelle and a signaling hub, relaying signals from a number of developmental pathways including the Sonic Hedgehog (Shh) pathway and regulating proper morphogenesis and patterning.^12^ Deletion of Kif3a - a gene essential for cilia formation - in the developing pancreas results in severe pancreatic defects, including acinar-to-ductal metaplasia, lipomatosis and fibrosis.^13^

Another interesting aspect of ciliary biology is that they are remarkably transient, yet critical, structures. Cilia are constructed from the docking of a primary centriole at the plasma membrane, which is then used as the foundation for microtubule extension. Hence, cell cycle plays a critical role in ciliary presence. An actively dividing cell cannot, therefore, maintain a mature cilium, as its centrioles are needed to coordinate chromosomal separation and cytokinesis. The critical role that motile and primary cilia play in development—the most active period of cellular proliferation—is therefore somewhat paradoxical. When and why do they appear where they do?

As the different cellular lineages organize with respect to one another in an organ, the cellular antenna that is a primary cilium is well situated to detect and coordinate short-range paracrine signals. Understanding which cells require primary or motile cilia can provide critical insight into these cross-lineage organizational relationships. However, detailed temporospatial analysis of ciliary dynamics has thus far been largely limited to the epithelium—likely motivated by the clinical relevance of motile ciliary defects to the innate immune functions of lung bronchial epithelium.^9,10,14^

In this study, we define the temporal presence or absence of cilia on different cell types in the developing mouse lung. Using immunofluorescent antibody staining, we find that pulmonary tissues display a high density of PC during early organogenesis. In addition, we identify the cells - epithelial, mesenchymal, or endothelial - that display cilia, throughout development and into postnatal life. As previously shown, we find that as development progresses, the epithelium transitions from mono-to multi-cilia proximally, while losing cilia in the distal lung. We find that endothelial cells (ECs) are almost completely devoid of PC at all stages of development examined, while mesenchymal cells initially express — but then steadily lose — their PC. A critical exception is the alveolar PDGFRα+ fibroblasts, which retain cilia into adulthood. By understanding what pulmonary cells display cilia (and when), we hope to gain insight into their functional roles during organogenesis, and the importance of cilia that are maintained into adulthood.

## Materials and methods

### Mice and embryo handling

Experiments were performed in accordance with protocols approved by the University of Texas Southwestern Medical Center (UTSW) Institutional Animal Care and Use Committee. Both male and female E10.5-E18.5 CD1 embryos and postnatal tissues were collected and dissected in phosphate-buffered saline (PBS). Animals were euthanized with CO_2_ followed by cervical dislocation, after which removal of lung tissue was performed directly. The *Pdgfrα-H2B-eGFP* mice were obtained from The Jackson Laboratory (Stock #007669).

### Dissection, sectioning and immunofluorescent analysis of lung sections

E12.5-E18.5 embryos were dissected and fixed in 4% paraformaldehyde (PFA) in PBS overnight at 4°C. Lungs of postnatal stages P0 and P7 were removed and placed directly into 4% PFA and fixed overnight. Lungs of postnatal stages P14 to P60 were inflated with intratracheal instillation of 4% PFA to 20 cm H_2_0 pressure. The trachea was tied off and the whole lung placed in 4% PFA for overnight fixation, after which individual lung lobes were dissected for further processing. Following dissection, lungs were processed for either cryosectioning or paraffin sectioning. In all immunofluorescent assays, coronal (frontal) sections of lungs were analyzed. Location of regions assessed are shown in all figures as boxed areas, showing region along the proximodistal axis of the lung, and whether the region is more central or peripheral. Analysis of epithelium, mesenchyme or endothelial cells is noted.

Following 3 washes in PBS, immunofluorescent stains were then performed as previously described (Daniel et al., 2018). In brief, tissues were dehydrated in an 25-50-75-100% ethanol series, cleared in xylene, and embedded in Paraplast (Leica) as previously described.^15^ Embedded tissues were sectioned at 10 μm thickness on a HistoCore MULTICUT semi-automated rotary microtome (Leica) onto SuperfrostPlus glass slides (Fisher). Sections were permeabilized for 10 min in 0.2% Triton X-100 in PBS, then subjected to antigen retrieval with R-buffer B in a 2100 Retriever (Electron Microscopy Sciences). Slides were blocked with CAS-block (Invitrogen), and primary antibody incubations were carried out at 4°C overnight (for antibody information and dilutions, see **Table S1**).

Alternatively, for imaging of thick lung sections, the fixed embryos or lung tissue were fixed as described above, washed three times in PBS, and then embedded in 4% agarose. These blocks were sectioned at 150 or 200 μm on a Leica VT1200 S vibratome. Vibratome sections were permeabilized with 0.3% Triton X-100 for 15 minutes, washed with PBS, blocked with CAS-block, and stained with primary antibodies overnight. The tissue was mounted on SuperfrostPlus glass slides with 0.25 mm deep iSpacer® double sided sticky wells.

For all immunofluorescent assays, coronal (frontal) sections of lungs were analyzed. Location of regions assessed are shown in all figures as boxed areas, showing region along the proximodistal axis of the lung, and whether the region is more central or peripheral. Unless otherwise specified, more distal regions of the lung parenchyma were chosen (see **Fig. 2G** schematics) at different stages of development (**Fig. 4**). Analysis of epithelium, mesenchyme or endothelial cells is noted. Endothelial cells of the larger arteries, arterioles, and veins were included when present, but they represent a small fraction of the total number of endothelial cells analyzed.

Images for all types of sections were obtained with either an A1R Nikon confocal microscope or a Nikon CSU-W1 dual camera inverted spinning disc confocal microscope.

Lung sections from *Pdgfrα-H2B-eGFP* mice were prepared for whole-mount staining as previously described.^16^ Briefly, the thick lung sections were permeabilized with PBS+1% Triton X-100 and blocked with PBS+0.3% Triton X-100+1% bovine serum albumin. Tissue sections were stained with primary antibodies for 48 hours, followed by overnight secondary antibody incubation. The samples were further cleared with Scale A2 for one week prior to imaging on a Leica Sp8 confocal microscope. Images were processed using ImageJ (FIJI) and the presented images are z-stack projections of 40 µm containing optical slices of 1 µm.

### Electron microscopy

Whole E12.5 embryos were fixed overnight in 4% PFA and sectioned at 100 μm. Sections were submitted to the UTSW Electron Microscopy core for further processing with standard protocols. Slices were imaged on a JEOL 1400+ transmission electron microscope.

### Cilia presence and length quantifications

Cilia were assessed using FIJI. To quantify the percentage of endothelial cells (ECs) harboring cilia, 100 ECs were assessed in each of 3 different fields of view (FOV). Visual sections of 5-10mm thickness had to be deconvoluted to distinguish individual cells. Ciliary length was measured utilizing both manual measurements and custom macros that trace the length of the ARL13b channel signal in three dimensions, beginning at the edge of the pericentrin (PCNT) signal and extending to the opposite end of the ARL13b signal. The measurements provided at this stage were taken from mesenchymal and epithelial cells at intervals along the length of the lung buds, both at the sides and the leading edge. Co-localization of ARL13b with IFT88 and acetylated alpha-tubulin was used to demonstrate that ARL13b staining reached the distal end of the cilium and support the accuracy of ciliary length measurements. For cell type-specific analysis, 3D analysis of cells that were positive for GM130 and ARL13B was necessary to establish the cell from which ciliary structures originated. Careful analysis of 10 μm sections using z-stacks of 0.2 or 0.5 μm slice depths was performed on all ECs, and all cases wherein an EC appeared to be ciliated were referred for 3D rendering and analysis (see **Fig. 2G** schematics) at different stages of development (**Fig. 4**). 3D reconstructions were generated using Imaris 10.0 (Oxford Instruments). Surfaces were drawn using intensity thresholding, followed by manual tracing and editing to clearly demarcate nuclear borders. All ECs (macro and microvascular) were included in this analysis. Models were generated for all ECs where the cilia’s originating cell could not be determined by visual sectioning of image Z-stacks obtained by confocal microscopy.

### Data analysis and visualization

Data were both analyzed and plotted in Graphpad PRISM 10.2.3. Statistical tests (indicated in the relevant methods sections and figure legends) utilized were the parametric unpaired Student’s t-test and the 2-Way ANOVA with Tukey’s multiple comparisons test. Statistical significance is noted as follows: ns = p>0.05; ∗ = 0.01<p<0.05; ∗∗ = 0.001<p<0.01; ∗∗∗ = 0.0001<p<0.001; ∗∗∗∗ = p<0.0001. Images were analyzed in FIJI. Figures and models were made using Microsoft Powerpoint, data were partially analyzed in Microsoft Excel and text was written in Microsoft Word.

### scRNAseq data analysis

Raw sequencing reads from previously published single-cell RNA-sequencing (scRNA-seq) data from previous studies^17–19^ were downloaded from the NIH GEO accessions GSE180822, GSE149563, and GSE165063, respectively (**Table S2**). Reads were mapped to the GRCm10 mouse reference genome using STAR-solo.^20^ Gene-by-cell counts matrices were subsequently ambient-RNA corrected, integrated, filtered, and projected using tools in the *scverse* toolchain^21^ as described in^16^. Cell type annotations were made manually by examining expression of known marker genes for each respective cell type within groups of cells within single/highly similar clusters.

## Results

### Primary cilia are present throughout multiple tissues in the early mouse embryo

To assess where primary cilia (PC) in the embryo might be positioned to play a role in tissue formation—beyond the well-known establishment of laterality by the cilia at the ventral node (E7.5)—we looked for presence of PC in a range of embryonic organs. We first stained sectioned mouse embryonic lungs using antibodies to ARLB13, which is a highly specific ciliary marker (**Fig. S1A**).^22^ To assess which specific cell types might harbor PC, we also stained for endothelium using antibodies to the membrane-bound glycoprotein endomucin (EMCN) and for epithelium using antibodies to the cell adhesion protein E-cadherin (ECAD/CDH1). Our results showed widespread presence of PC throughout embryonic organs at embryonic day 10.5 (E10.5, **Fig. S1A-F**). We noted a high frequency of PC not only in the epithelium of tissues like lung (**Fig. S1A**) and gut (**Fig. S1B**), but also throughout the surrounding mesenchyme (asterisks). ARL13B+ structures could be observed scattered throughout all organs and tissues examined.

To further evaluate distribution, we made a global quantification of total cilia and total number of cells in multiple organs at E10.5. We counted the number of cells observed (using DAPI to identify nuclei of single cells) that displayed an associated ARL13B+ structure, as well as the total number of cells that did not. Overall, we found a high PC-to-cell ratio in multiple organs, particularly the lung (75%), gut (76%), and neural tube (71%), while tissues such as the liver and heart exhibited fewer cilia-bearing cells (**Fig. S1G).** Overall, these observations suggest a role for cilia during mammalian organogenesis, as they are widely distributed in different organs early during development, including lung.

### Primary cilia are present in multiple cell types in the embryonic lung

We next assess cilia in the budding lung more closely. Examination of E11 lungs revealed that cilia positive for ARL13B staining were widely distributed throughout the lung, and present in both proximal and distal regions (**Fig. 1A**). We next sought to determine the frequency of cilia on the main cell types - epithelial, mesenchymal, endothelial (**Fig. 1B-D**).

**Fig. 1.**
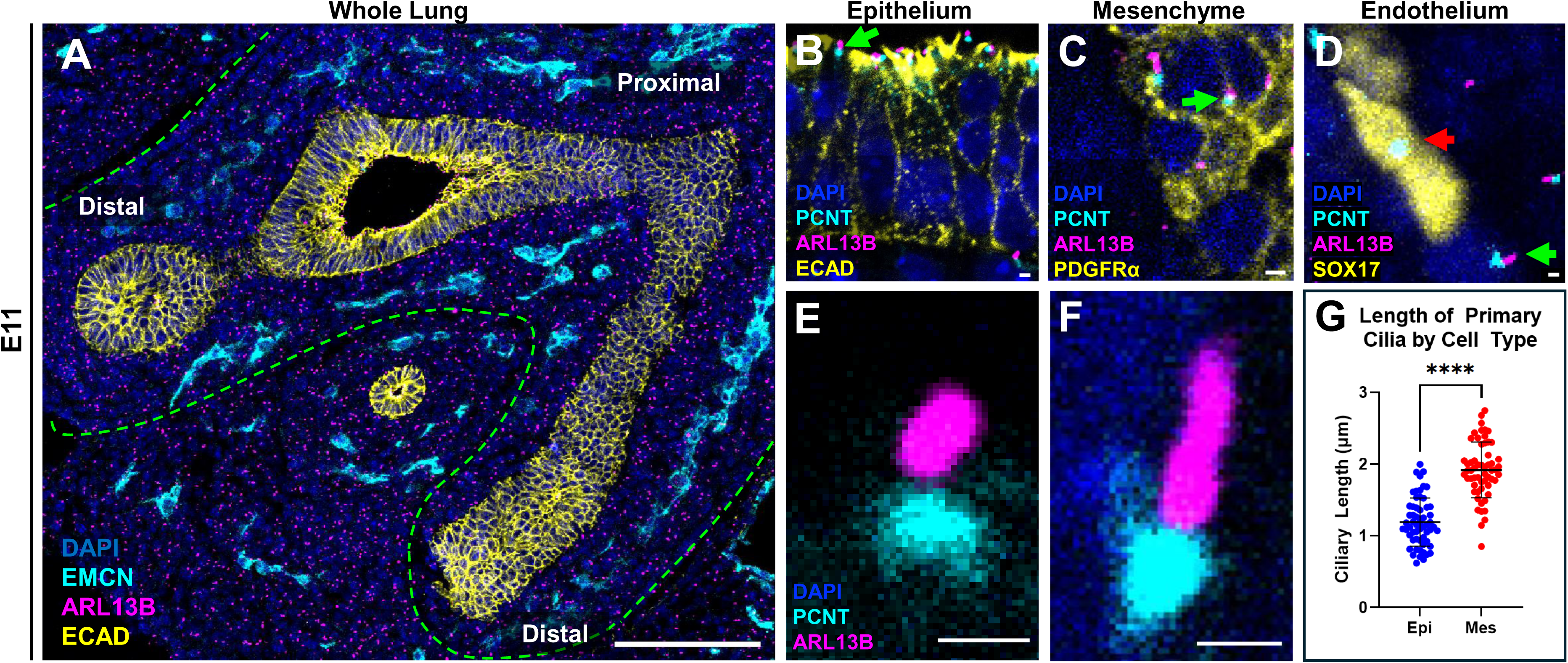
Presence of primary cilia of distinct lengths in lung epithelial and mesenchymal cells. At E11, immunofluorescent staining of developing lung tissue for PC with ARL13B **(A)**. Higher magnification views of co-staining for PCNT and ARL13B (green arrows) in epithelial (ECAD, **B**), mesenchymal (PDGFRα+, **C**), and endothelial (SOX17+, **D**) cells. Note presence of PCNT, but absence of ARL13B in the EC (red arrow) in **(D)**. Adjacent non-endothelial cells display structures that are positive for both PCNT and ARL13B (green arrow). The cilia expressed by epithelial cells **(E)** are significantly shorter than those expressed by mesenchymal cells **(F)**. Mean epithelial ciliary length was 1.19 µm, while mean mesenchymal ciliary length was 1.92 µm **(G)**. Cilia in E, F were assessed in distal lung sections. Scale bars are 100 µm (A) and 1 µm (B-F).

Focusing first on the pseudostratified epithelium of the distal lung bud, we found that each epithelial cell was associated with a single PC (**Fig. 1B**). The presence of a single cilium in each cell was confirmed by co-staining for ARL13B and pericentrin (PCNT), which localizes to the base of each PC. The columnar organization of the epithelium made it straightforward to identify cell of origin for each cilia. However, the more complicated organization of both mesenchyme and endothelial cells (ECs) in distal portions of the lung was more challenging. Accuracy was further limited by a high density of embryonic tissue cells with a high nuclear-to-cytoplasmic ratio. To precisely assess the cell types that harbored cilia in these tissues, we sought to establish the polarity of each cilium by staining for cilia-associated structures found at the proximal end of the structure. We did this by defining the 1:1 cilium:cell ratio using examination of cells in 3 dimensions, using Maximum Intensity Projection (MIP) reconstructions of Z-resolution (0.5 mm). This facilitated identification of cilia in mesenchymal cells (**Fig. 1C**). Indeed, staining for the mesenchymal marker PDGFR*α*, allowed us to ascertain that most mesenchymal cells within the lung bud displayed single PC.

We next asked whether ECs also harbored cilia (**Fig. 1D**). While PC in epithelium and mesenchymal cells were pervasive and easily identifiable, it was more difficult to find PC on ECs. Sox17+ ECs mostly exhibited PCNT staining with no associated ARL13B signal (red arrow), while cells in adjacent mesenchyme displayed clear colocalization of PCNT and ARLB13 positivity (green arrow) (**Fig. 1D**). Together, these data show that the PC is differentially present among multiple cell types during early lung organogenesis.

Of note, the combined ARL13B and PCNT stain also allowed for evaluation of individual cilia length (**Fig. 1E, F**). We found that PC in the epithelium (**Fig. 1E**) displayed shorter cilia than those in the mesenchyme (**Fig. 1F, G**). Given that ciliopathies present with shorter cilia, this differential length suggests that the epithelial primary cilia may have different signaling functions than those in the mesenchyme at this stage.^23^

To confirm that ARL13B marked bona fide cilia, we carried out costains for other known markers of PC, including Ift88, γ-tubulin and acetylated α-tubulin (**Fig. 2**)^24,25^. ARL13B does indeed label cilia that were also positive for these additional markers, at different stages and throughout the lung. To further confirm cilia in the mesenchyme at this stage, we performed Transmission Electron Microscopy (TEM) on E12.5 lungs (**Fig. S2**), which demonstrated PC located in a deep pocket on mesenchymal cells (**Fig. S2B**), with basal body and axoneme clearly visible (**Fig. S2B, inset**).

**Fig. 2.**
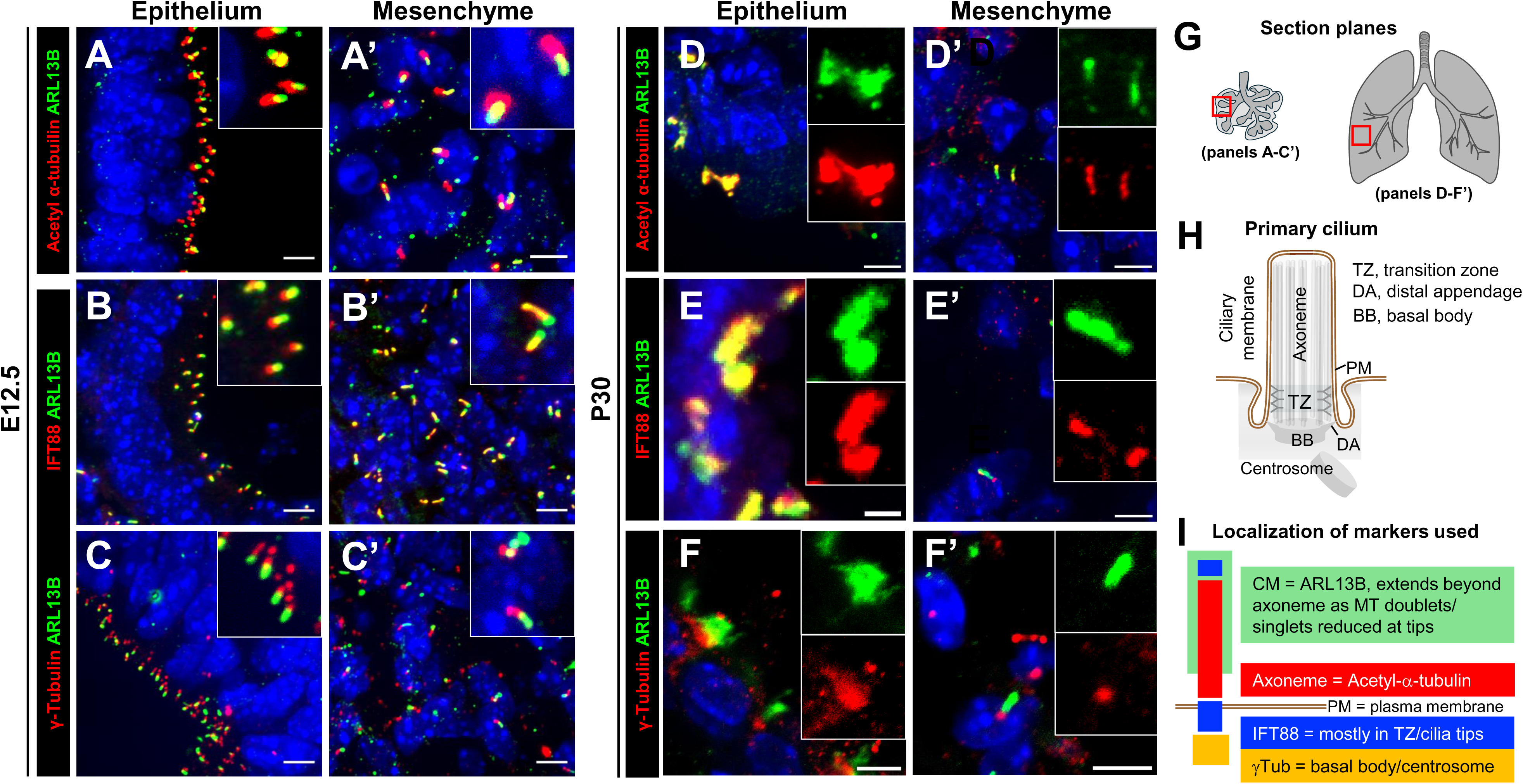
Confirmation of primary cilia identification with multiple immunofluorescent markers. Immunostaining of lung tissue at E12.5 (**A-C’**) and P30 (**D-F’**). (A). In all panels, ARL13B is shown in green, while other ciliary markers including acetyl-a tubulin (**A, A’, D, D’**), IFT88 (**B, B’, E, E’**), and g-tubulin (**C, C’, F, F’**) are in red. (G) E12.5 images were selected from the distal lung bud epithelium and surrounding mesenchyme. P30 images were selected from the distal bronchioles (to show motile cilia on multiciliated epithelial cells) and the surrounding distal alveolar space. (H) Structure of a primary cilium and location of different substructures. (I) Localization of immunofluorescent markers used, showing proximodistal specific locations. Scale bar is 5 μm.

### Cell type specific evaluation of cilia requires 3D analysis

The difficulty in clearly identifying endothelial cilia (**Fig. 1D**) prompted us to assess additional ciliary markers to more clearly delineate the cell of origin for each PC. We therefore used GM130 to visualize the Golgi. Similar to PCNT, the Golgi is always associated with the base of the cilia (**Fig. 3A,B**), but it additionally wraps around the nucleus of the cell of PC origin (yellow bracket). This shared contour allows for a more confident assignment of an individual cilium to its cell of origin.

**Fig. 3.**
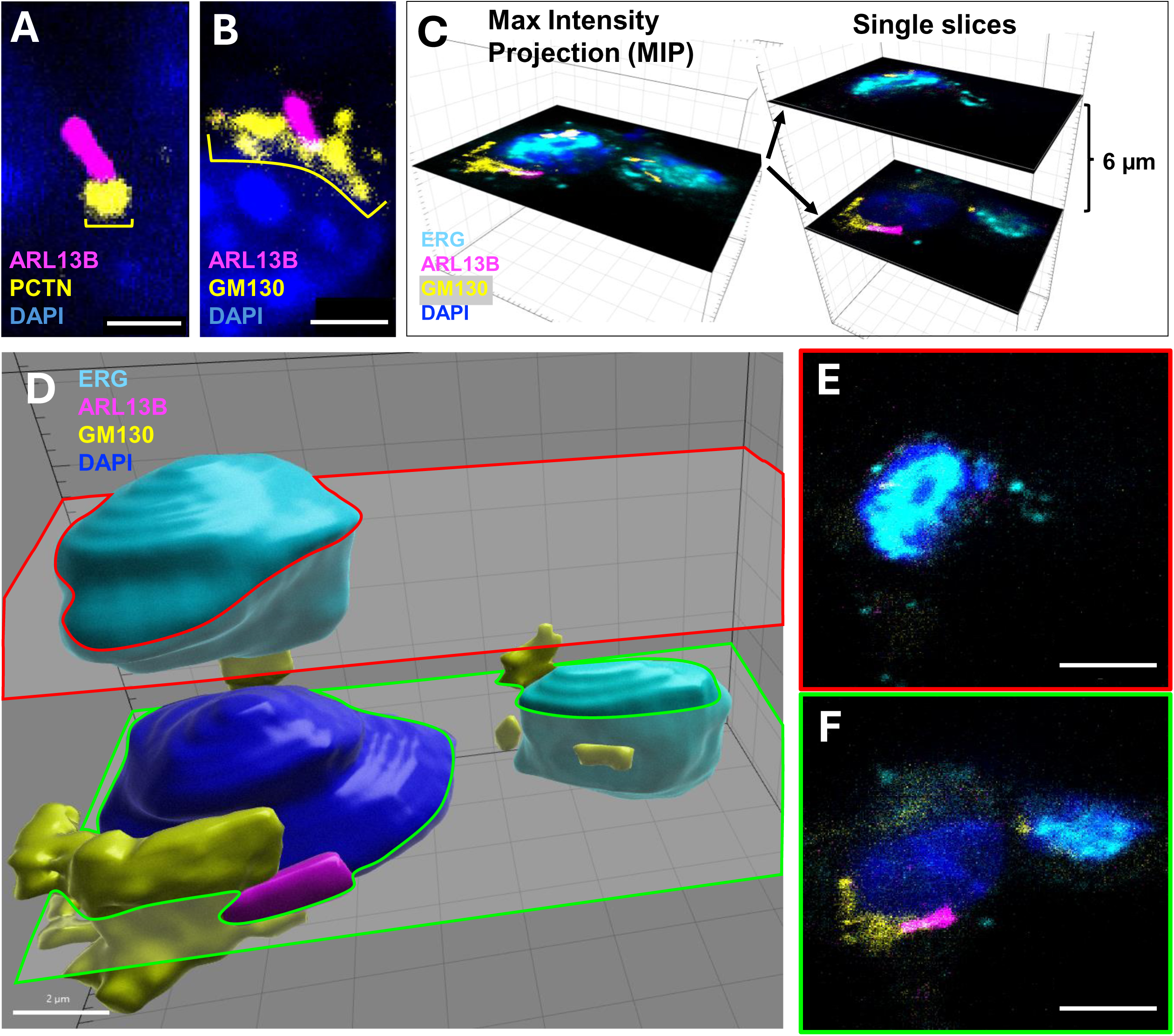
Identifying ciliary cell of origin at the cellular level in embryonic tissues requires 3D analysis. Pericentrin (PCNT) marks the basal body of cilia **(A)**. Golgi (GM130) also marks the base of a PC but provides contour to match the nucleus of the cell that expresses the cilium **(B)**. The contour of GM130 allows for clear differentiation of cell-of-origin, while high resolution (0.2-0.5 µm slices) confocal microscopy **(C)** and 3D modeling with Imaris **(D)** can overcome both high cell density and overlapping cells of different types **(E, F)**. Scale bar is 5 µm.

To assess possible endothelial cilia, we additionally used the endothelial nuclear marker ERG, which is expressed in most ECs in mid-gestation stage lung tissue,^26^ in combination with ARL13B and GM130 (**Fig. 3C**). However, we realized that for ECs, using maximum intensity projection (MIPs) still yielded artifactual data, as identifying cell of origin for a single cilium depended on the precise z plane examined. As this method (MIP) compresses multiple visual slices, it often inaccurately showed ECs has having cilia. We therefore turned to analyzing ECs using 3 dimensional reconstructions (**Fig. 3A’-D’**). Single slices helped provide additional granularity (**Fig. 3E, F**) and allowed us to analyze cell of origin for each PC within the interstitium of the lung parenchyma. The methodology aided in the rigorous identification of embryonic lung EC and the possible presence of cilia. Using this approach, as well as careful analysis of 3D confocal Z-stacks, we found that mouse embryonic lung EC do not exhibit ciliary structures.

### Lack of primary cilia in the pulmonary endothelium during development

Using the methods developed above, we assessed ciliary status in ECs throughout lung formation. We carried out 3D immunofluorescent staining of ARL13B in combination with SOX17, an EC nuclear marker and GM130, for identification of Golgi. We focused on the distal lung, during both pre- and post-natal development.

Similar to our observations at E11 (**Fig. 1D**), we found that ECs examined in 3D lacked PC at all developmental timepoints (**Fig. 4**). Sectioned embryonic lung from E12.5 (**Fig. 4A**), E16.5 (**Fig. 4B**) and P0 (**Fig.4C**), as well as adult tissue P30 (**Fig. 4D**) was stained for ARL13B, GM130 and SOX17. We assessed ECs located within the stroma of distal alveoli as we did in **Fig. 2 (see schematic in 2G)**. 3D renderings showed that SOX17+ nuclei were devoid of ciliary structures. Using ARL13B combined with GM130 allowed determination of the cell of ciliary origin, and we showed that SOX17+ nuclei were not associated with PC (**Fig. 4A’-D’**). We counted 1333 nuclei in 12 fields of view (FOV) in the lung periphery, and we observed no association of ARL13B/GM130 with SOX17+ nuclei (**Fig. 4E, F**). Complete absence of PC was noted in all embryonic ECs as well as mature ECs after the completion of alveologenesis (after P30). We did not attempt to differentiate between alveolar (aCap) or general (gCap) capillary cells in this study, as both cell types express ERG^27,28^ and no ciliated ERG+ (or SOX17+) cells could be found at later time points (**Fig. 4C-D**).

**Figure 4.**
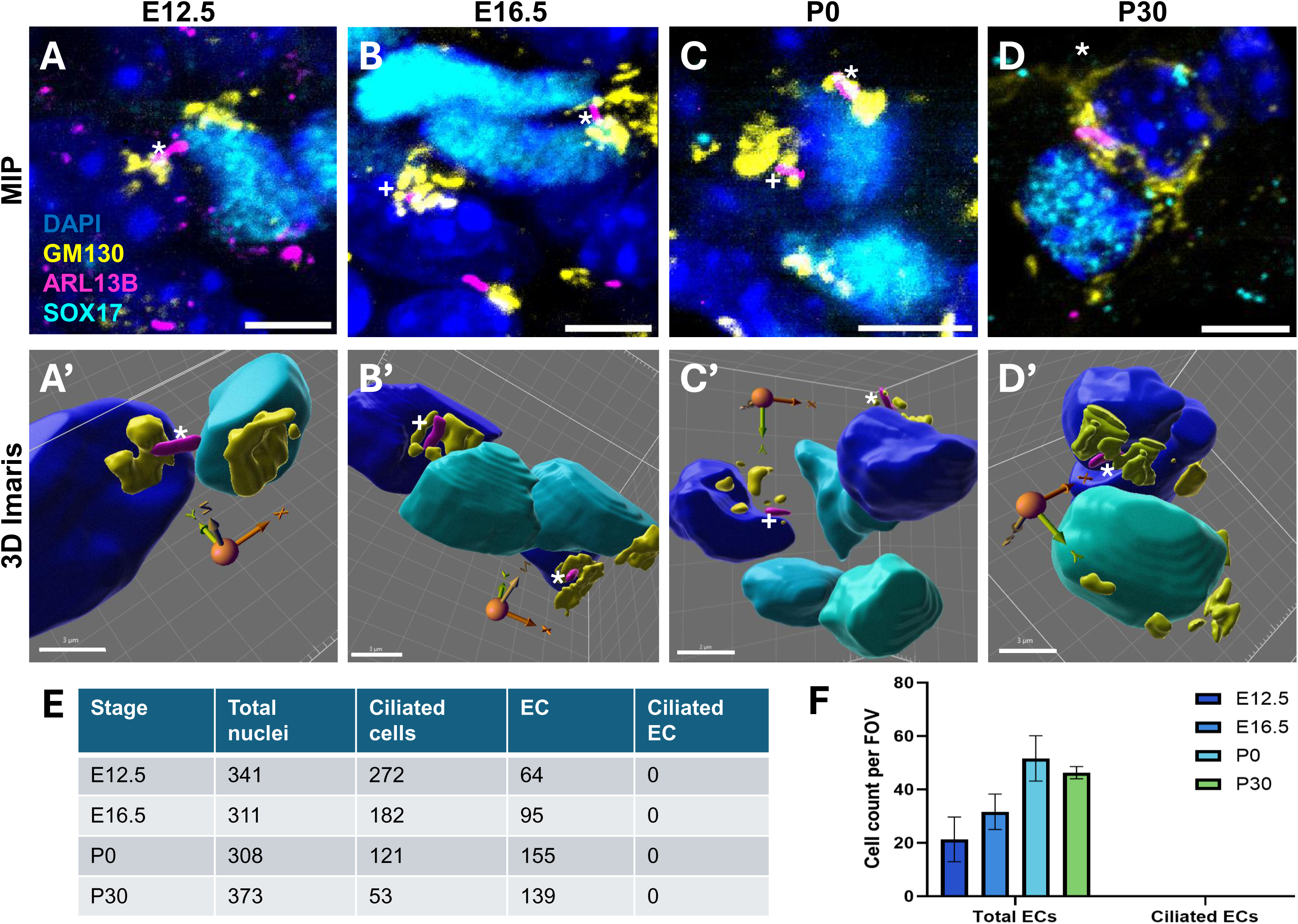
Pulmonary endothelial cells lack primary cilia throughout development. Immunofluorescent staining of 10 µm slices of PFA-fixed, paraffin-embedded mouse lung tissue at the indicated developmental time points **(A-D)**, followed by 3D rendering of the nuclei **(A’-D’)** to determine each cilia’s cell of origin. 3D models of nuclei, golgi, and cilia color the match the IF stains above. X and Y correspond to the X and Y plane of the above image, with model rotated to highlight each cilia’s correct relationship to the nucleus of its cell of origin. * and + identify the same cilia in each matching image set (**A, A’** for example). Quantification of total cells analyzed **(E)** reveal no ciliated endothelial cells among those analyzed at E12., E16.5, P0, or P30, while the mean EC and ciliated EC counts per FOV analyzed are shown in **(F)**. Sections taken in regions similar those in **Fig. 2G**. Scale bar is 5 µm.

Because it has long been known that ciliary structure is dependent on the state of the cell cycle,^29^ we next examined whether perhaps the apparent lack of EC cilia was due to ECs actively cycling. To address this point, we co-stained ECs with the endothelial marker ERG and the marker of cell proliferation Ki67 (which shows no staining in G0, punctate nuclear staining in G1 and G2, and strong nuclear staining in M phase). Actively dividing cells were easily identified (**Fig 5, A-D’**, white arrows). We found that while a small percentage of ECs at each stage examined were actively proliferating (**Fig. 5E**), Erg+ cells that were not cycling also did not exhibit ARL13B+ structures (**Fig. 5, A-D’,** green arrows). These findings indicate that the lack of endothelial PC is not likely representative of regression of cilia due to cell division.

**Fig. 5.**
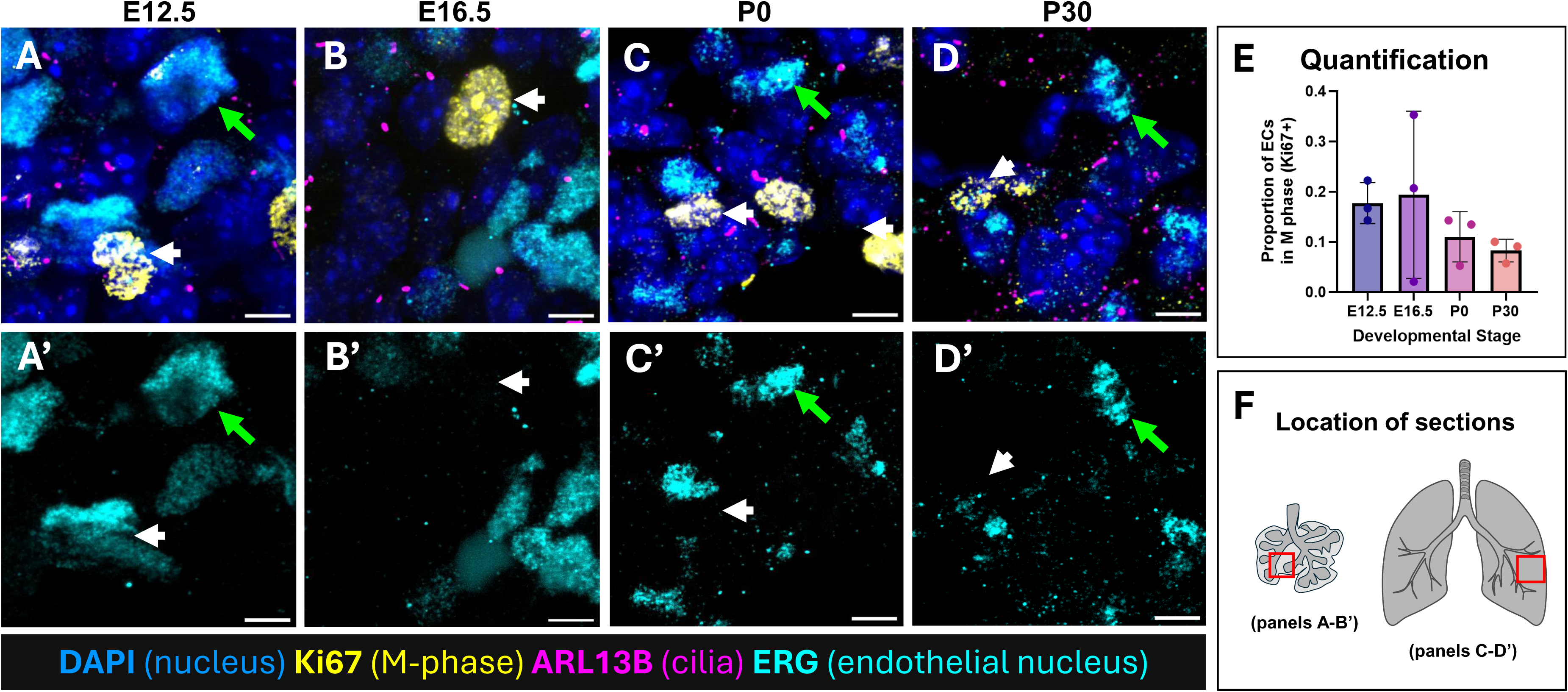
Absence of cilia in endothelial cells is not due to cilia regression during mitosis. Images showing lung endothelial cells (Erg, cyan) co-stained for the proliferation marker Ki67 (yellow) **(A-D)**. Panels showing Erg only staining **(A’-D’)**. Note absence of ARL13B staining in dividing cells that are positive for Ki67 (white arrows). Conversely, note that numerous Erg+, but Ki67 negative cells (green arrows) do not display ARL13B+ cilia. **(E)** Quantification of ECs demonstrating Ki67 positivity. 1333 cells were counted. **(F)** Location of section for sections in **(A-D’)**.

While we found the distal lung endothelium was devoid of PC, ECs in larger vessels did indeed display PC, as previously reported. Whole mount staining of *en face* dissected branch pulmonary arteries revealed that the more proximal, large branch pulmonary arteries did contain ciliated ECs in the adult mouse (**Fig 6A, C**). Similarly, we were able to identify PC in ECs of the aortic arch (lesser curvature) (**Fig. 6B, D**). These findings are in keeping with previous work reporting presence of PC in the aorta—specifically, along the lesser curvature of the arch, where turbulent flow is present.^30^ We confirmed this observation using TEM (**Fig. S3**), which revealed absence of PC in regions of laminar flow (descending aorta) (**Fig. S3A**). Regions of disturbed flow, like the lesser curvature of the aortic arch, showed some occasional PC (**Fig. S3B, C**). Hence, we were able to find some examples of endothelial PC, albeit in adult EC.

**Fig. 6.**
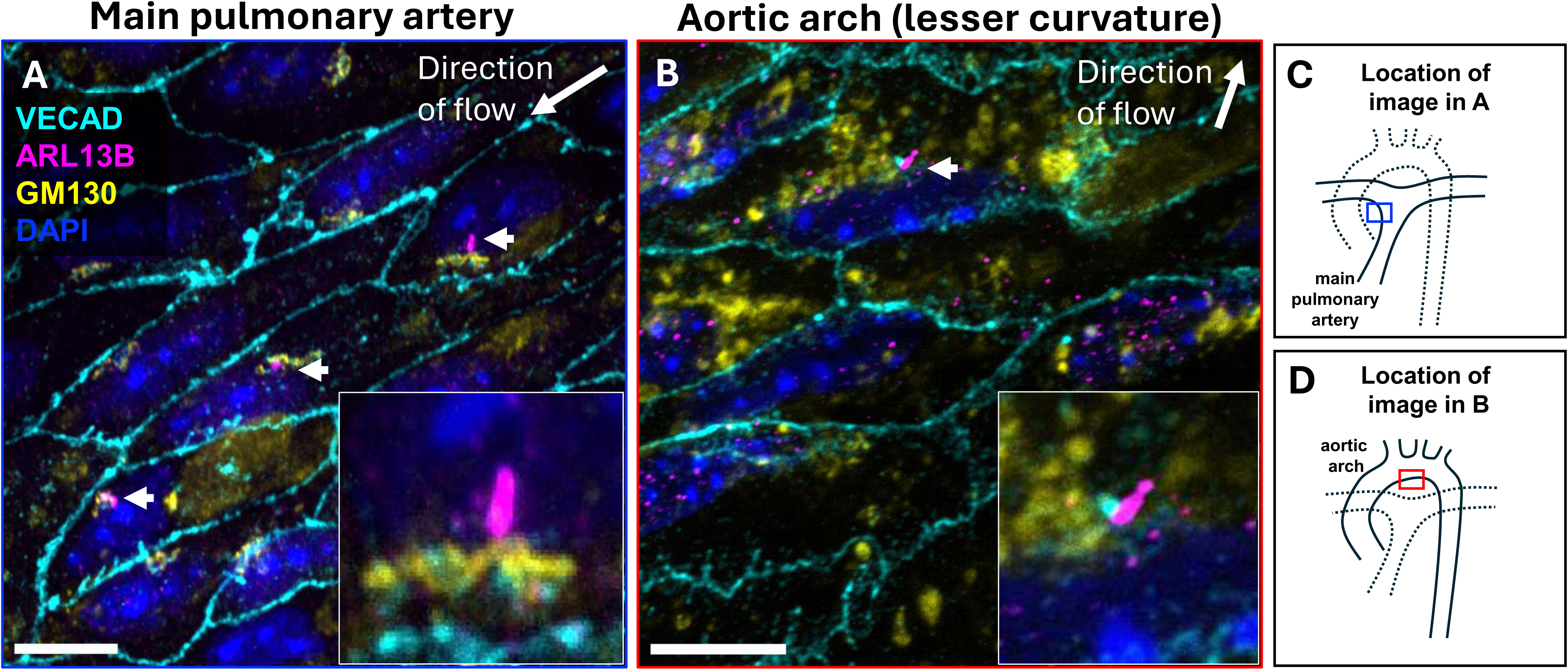
Presence of cilia in the large pulmonary arteries and aorta. Aorta and pulmonary artery of mice (n=3, P60) were perfused and fixed with PFA prior to *en face* staining for EC borders (VECAD, cyan) and primary cilia (ARL13B, pink), with Golgi (GM130, yellow) to clarify cell of origin. Multiple cilia were found in both pulmonary artery **(A)** and the lesser curvature of the aorta **(B)**. Schematics showing location of stained ECs (boxed areas) in main pulmonary artery (**C**, blue box) and in the aortic arch (**D**, red box). Scale bar is 10 µm.

This work identifies a proximal-distal gradient of endothelial PC in large vessels of the adult mouse, with select subsets of ECs in the large arteries displaying PC, whereas EC of the distal capillaries do not.

### Proximal-distal gradient of ciliary fate in developing pulmonary epithelium

In the adult lung, the presence of PC has only been reported in the airway smooth muscle,^31^ while the presence of multiciliated epithelial cells in the large airways is well known.^11^ Conversely, the highly specialized AT1 and AT2 cells of the mature alveolus lack cilia of any kind.^32,33^ The dynamics of how these different regions of the lung acquire different types of cilia has not been examined.

We sought to assess epithelial ciliary structures along the proximal-distal axis of the lung epithelium (**Fig. S4A**). Using immunofluorescence for FOXJ1 (a master regulator of multiciliation), we saw that in the mouse lung, the onset of multiciliation in the upper airways has occurred by at least E16.5 (**Fig. S4B**). As previously reported, not every cell in the adult upper airway is ciliated - goblet, club, basal, and tuft cells all lack cilia ^34^. We also noted the absence of PC in non-FOXJ1-expressing cells in the upper airways (**Fig. S4A’**). In distal regions of the lung, we found that epithelial cells at E16.5 did not express FOXJ1, but did display primary cilia (**Fig. S4B’**). As pulmonary development and alveologenesis progress, the distal epithelial cells become AT1 and AT2 cells, both of which lack cilia in the mature animal (**Fig. S4D’**). At E16.5, the developing epithelium thus appeared to have segregated into proximal FOXJ1+ multiciliary epithelium, and distal FOXJ1-single ciliary epithelium, depending on their location in the proximal-distal axis of the organ. Proximal expression of FOXJ1 in the developing large airways is directly correlated with apical clusters of ARL13B signal indicating likely multicilia (**Fig. S4**).

This pattern of proximal multiciliation and distal single cilia continued to be observed in postnatal lungs. In P0 lungs, we confirmed multiciliated cells in proximal airways using spinning disk microscopy. Using staining for PCNT, we showed that tufts of ARLB13+ structures were anchored by PCNT+ array of docked centrioles (**Fig. S4C**). At P60, these structures were clear in the proximal columnar epithelial cells of the upper airway which displayed many cilia per cell (**Fig. S4D**). By contrast, distal epithelial cells displayed single ARLB13 and PCNT positive structures both at P0 (**Fig. S4C’**) and P60 (**Fig. S4D’**) indicating presence of single PC.

### Maintenance of primary cilia in PDGFRα+ alveolar fibroblasts

Finally, because PC are observed in early lung mesenchyme, we assessed the presence of PC in the adult derivatives of the mesenchymal cell population, with a focus on the distal alveolae. Using DNAH11 as a reliable marker of motile cilia,^24^ we confirmed the presence of DNAH11 in motile epithelial cilia (**Fig. S5A**) in order to show its absence in the primary cilia identified in the alveolar region (**Fig. S5B**). In this region, PDGFR*α*+ cells comprise the majority of the lung parenchyma and are critical units of lung function.^35^ Given the complex three-dimensional architecture of the mature alveolus, we examined markers specific of all major cell types to assess which cells exhibit a primary cilium at the alveolar level. We found that while PDGFR*α* expressing cells exhibited single PC (**Fig. 7H**), neither pericytes (PDGFRβ+) (**Fig. 7B**), aCAP cells (CA4+) (**Fig. 7C**), ECs (SOX17+) (**Fig. 7D**), AT1 cells (HOPX+) (**Fig. 7E**), AT2 cells (LAMP3+) (**Fig. 7E**), nor lymphatic ECs (PROX1+) (**Fig. 7G**), displayed PC. Interestingly, the mesothelium also displayed primary cilia (**Fig. S6**). In sum, the mesenchymal cell population diversifies greatly as development progresses, but the PC presence appears to be primarily maintained in adulthood by PDGFR*α*+ alveolar fibroblasts.

**Fig. 7.**
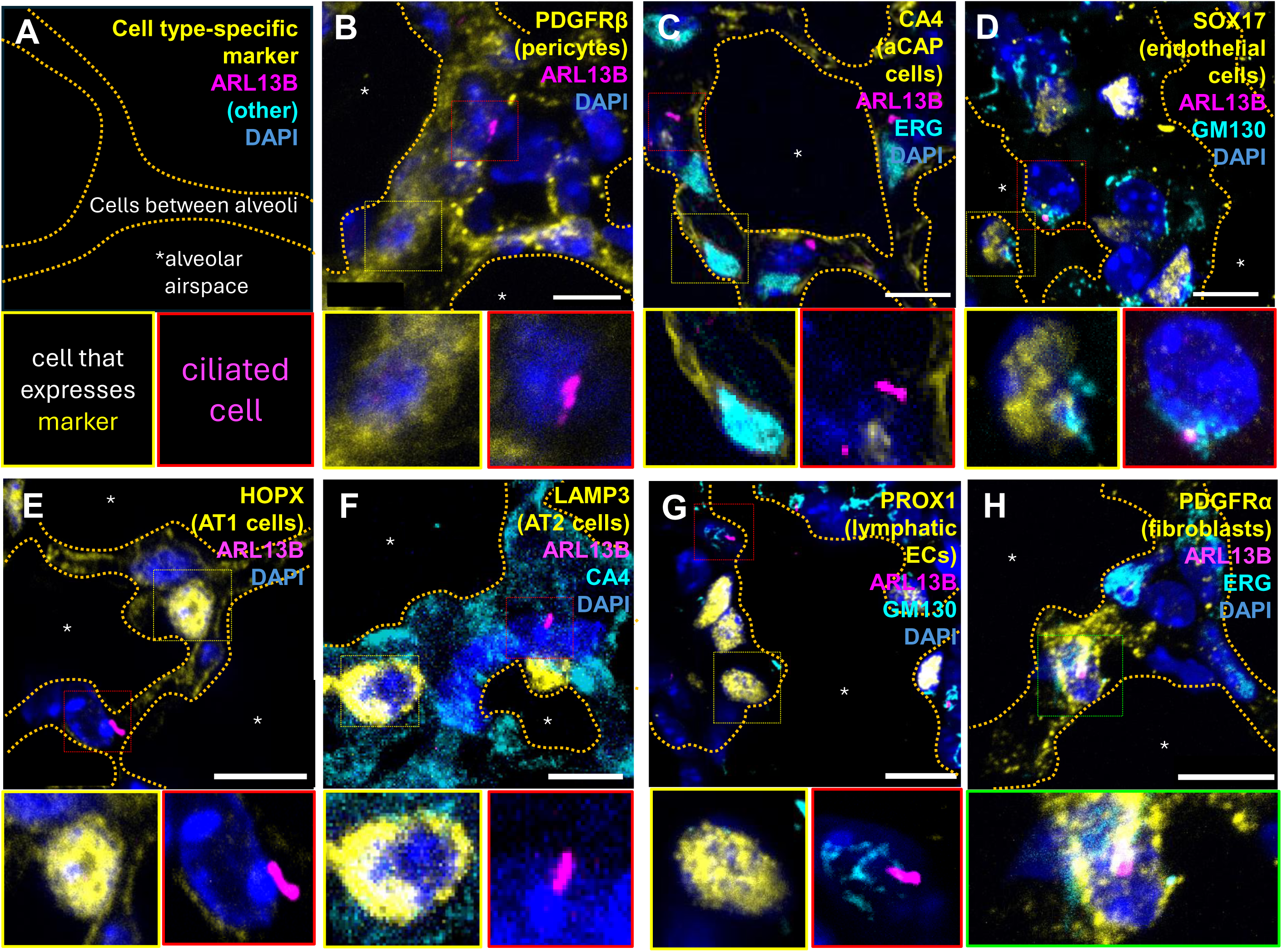
Maintenance of primary cilia by PDGFR*α*+ myofibroblasts in the adult at P30. Immunofluorescent analysis of adult mouse distal lung to identify cell types that harbor cilia. Figure organization is shown in **(A)**, with alveolar spaces outlined in orange and indicated by asterisks. In each image, yellow staining marks the cell type of interest. The yellow inset shows cells positive for cell type specific marker as indicated, while the red inset shows a ciliated cell. Immunofluorescent analysis revealed that pericytes **(B)**, alveolar capillary (aCap) cells **(C)**, ECs **(D)**, AT1 cells **(E)**, AT2 cells **(F)**, and lymphatic ECs **(G)** all lack PC. Co-localization of PC with is observed only with PDGFRα+ fibroblasts **(H)**. Scale bar is 10 µm.

### Temporal regulation of mesenchymal ciliogenesis as analyzed by scRNAseq

To further gain insight into the specific mesenchymal cell types that might harbor PC in the mouse lung, we carried out analysis of previously published scRNAseq datasets^17–19^ (**Fig. 8A**). Focusing on the mesenchymal subset, UMAP analysis was able to delineate all the expected cell types, including airway smooth muscle cells, fibroblasts and secondary crest myofibroblasts (SCMFs) (**Fig. 8B**). We also resolved cell types by developmental stage (E12, E17, P7, P15, P42) (**Fig. 8C**).

**Fig. 8.**
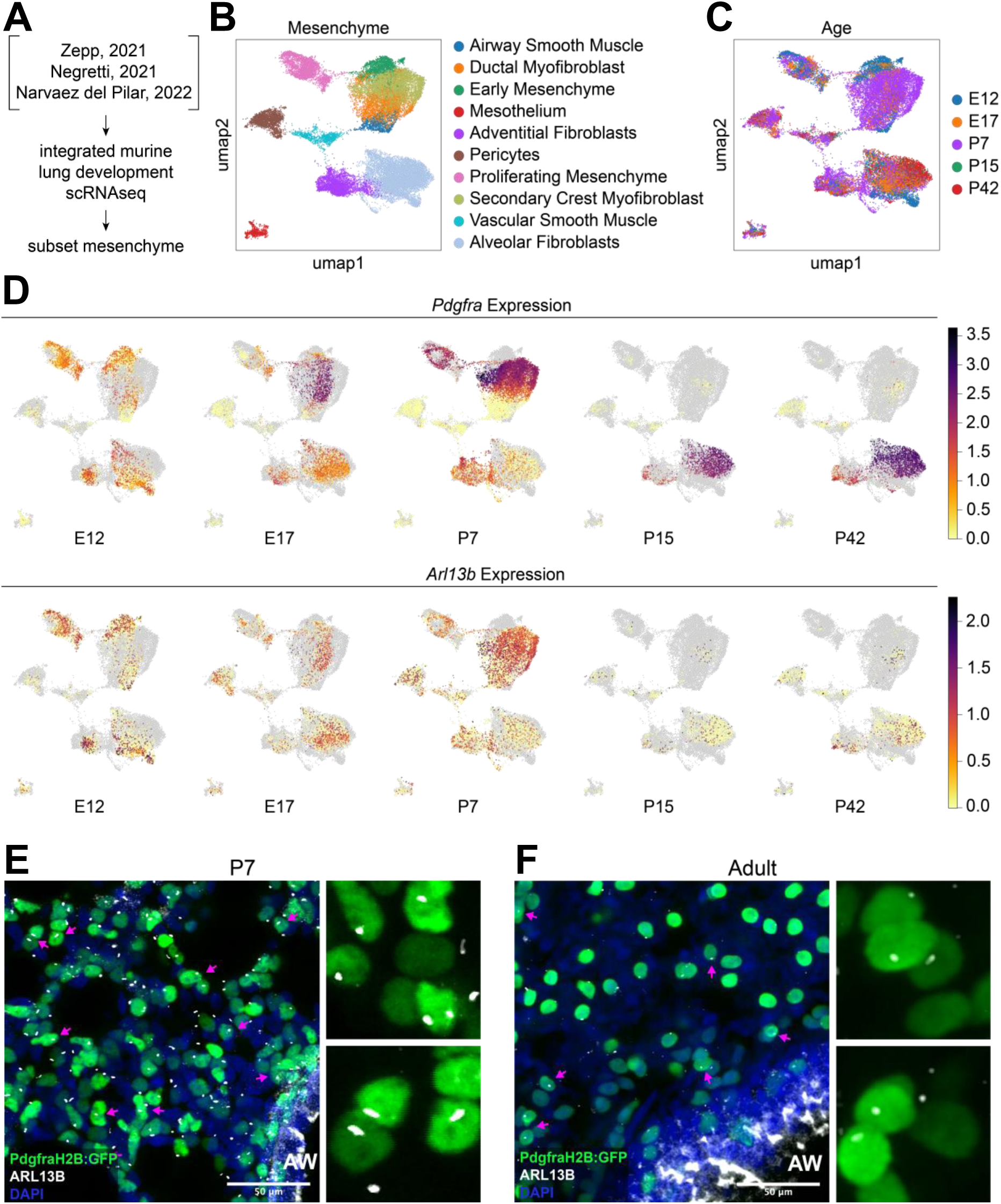
Mesenchymal *Arl13b* expression can be found in *Pdgfrα-*expressing fibroblasts throughout lung development. **(A)** Previously published lung development scRNAseq datasets were integrated and the mesenchyme was included as a subset. **(B)** UMAP of mesenchymal cell types **(C)** UMAP of developmental timepoints. **(D)** Feature plots show expression of *Pdgfrα* and *Arl13b* in the lung mesenchyme across developmental timepoints. Whole-mount images (40µm z-stacks) from P7 **(E)** and adult **(F)** *PdgfrαH2B:GFP* reporter mice shows nuclear ARL13B expression in *Pdgfrα*-expressing fibroblasts (magenta arrows). Scale bar is 50 µm.

Using the single transcript *Arl13b* to assess correlation between RNAseq and our immunofluorescent analysis of mesenchymal cells, we examined clusters that represented various subsets of stromal cells, including early mesenchyme, fibroblasts and various types of smooth muscle and myofibroblasts. Examining embryonic stages (E12 and E17), we found that *Pdgfrα* was highly transcribed in adventitial and alveolar fibroblasts, early proliferating mesenchyme and ductal fibroblasts (**Fig. 8D**). Postnatally, *Pdgfrα* expression peaked in proliferating and SCMF mesenchymal populations, as well as in adventitial fibroblasts. After P15, elevated expression was observed mainly in alveolar fibroblasts. Strikingly, these were all the same populations that displayed high levels of *Arl13b* transcripts, reflecting our immunofluorescent findings.

To assess whether scRNAseq data would confirm the lack of PC in the endothelium, we created an endothelial-specific subset from the same data (**Fig. S7**). This analysis revealed sporadic cells that expressed *Arl13b*, with a higher number of *Arl13b*+ cells in the gCap cells of the older (P42) animals (**Fig. S7D**). We then asked if these *Arl13b*+ cells were also positive for *Ift88* and *Tubg1*, as we had seen clearly by immunofluorescence. By creating a sequential subset of *Arl13b, Ift88,* and *Tugb1*, we were able to investigate the ability of a limited gene set to identify ciliated cells (**Fig. S8A**). Unfortunately, while the cilia themselves clearly contain the protein, the transcripts were not all present in similar abundance within the same cells. The “ciliated” subgroup of epithelial cells was the only group for whom even co-expression of two different transcripts (*Arl13b*+ and *Ift88*) did not reduce the proportion of positive cells to a negligible value (**Fig. S8B**). In fact, even the epithelial cell population as a whole was reduced from 10,901 cells to only 22 cells that co-expressed *Arl13b, Ift88, and Tubg1*.

We then broadened our examination further to include all subsets—endothelial, mesenchymal, epithelial, and immune cells (**Fig. S9**). Beyond the findings in the mesenchymal subsets, we noted similar patterns of *Arl13b*+*, Ift88, and Tugb1* expression in epithelial cells, secondary crest myofibroblasts, and alveolar fibroblasts (**Fig. S9E**). These data reveal the overall patterns but do not facilitate identification of ciliated cells for downstream pathway analysis.

Given the clear overlap of *Arlb13b* expression with that of *Pdgfrα*, and the immunofluorescent data from Fig. 7, we asked whether the *Pdgfrα* population contributes to cells bearing PC in the postnatal lung mesenchyme. Using a *PdgfrαH2B:GFP* mouse line, we confirmed that the *Pdgfrα*-positive cells indeed give rise to cells displaying clear ARLB13+ structures at both P7 (**Fig. 8E**) and adulthood (**Fig. 8F**).

Together these data reveal a dynamic spatiotemporal arrangement of primary and motile cilia throughout the development of the murine lung **(Fig. 9).** The unique absence of primary cilia from ECs highlights the cell- and context-specific signaling in which primary cilia participate. We confirm the differential multiciliation and loss of epithelial cilia, and identify a population of PDGFRα+ alveolar fibroblasts that maintain primary cilia into adulthood.

**Fig. 9.**
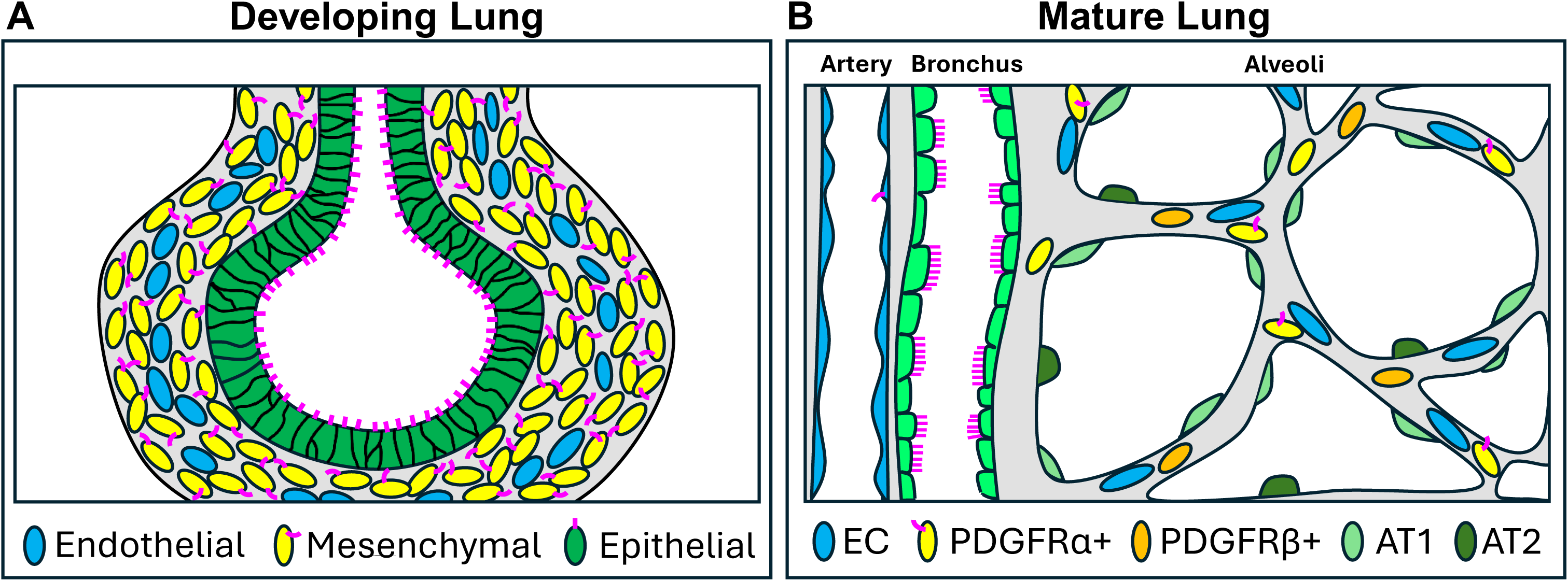
Summary of dynamics of ciliary presence in the developing and mature lung. **(A)** During embryogenesis, PC are observed along the apical surface of the lung epithelial cells (distal epithelium in dark green, and proximal bronchial epithelium in light green). In addition, cilia are observed on most mesenchymal (yellow), but not endothelial (blue) cells. **(B)** In the adult, PC are exclusively observed on PDGFRα+ cells within the stroma of the alveoli, but not PDGFRβ+ (orange), AT1 (light green) or AT2 (dark green). PC are not observed in the ECs (blue) of the aCaps. Some PC are found in the aorta and large pulmonary arteries, and multiciliated cells are found in epithelium (white) of the bronchus.

## Discussion

Here, we report the dynamic appearance of cilia in epithelial, mesenchymal and endothelial cell (EC) types within the developing mouse lung. We use immunofluorescent staining of markers for the ciliary protein ARL13B, as well as co-staining with pericentrin (PCNT) and GM130, to identify cell of origin of primary cilia (PC). We confirm that these ARLB13+ structures are cilia using other known ciliary markers. We examine forming lung tissues from E10.5, throughout development, and into adulthood. We find that contrary to our initial expectations, ECs within peripheral lung tissues never display PC. Endothelial PC are exclusively observed in large vessels, such as the pulmonary arteries, and the midline thoracic dorsal aortae. Conversely, cilia are found dynamically in epithelial and mesenchymal cell types. We find that bronchial epithelial cells display multicilia primarily late during development and into postnatal stages, while distal epithelial cells never harbor PC. Intriguingly, we observe PC within the pulmonary stroma, primarily on PDGFR*α*+ cells. We do not observe PC on numerous other cell types, including alveolar capillary (aCAP) cells, lymphatic ECs, AT1 and AT2 cells, nor pericytes. To our knowledge, this is the first high resolution characterization of cilia within the developing lung. These findings underscore the dynamic and cell specific presence of PC in lung and suggest important signaling pathways that are active within a subpopulation of the stroma.

### What role do cilia play in pulmonary organogenesis?

Our study aims to bridge two key findings well established in the literature — paracrine signaling and cilia signal transduction — and map these events at a cellular level in the developing lung. First, multiple studies have demonstrated the critical role of hedgehog, Wnt, and PDGF signaling in the normal development of the lung, as they coordinate cellular outgrowth and differentiation to form a branched and vascularized epithelial structure.^36,37^ Second, numerous studies have identified the ciliary localization of hedgehog Wnt, and PDGF signaling receptors and downstream effectors, and the role cilia in effective signal amplification.^38^ The role of this remarkable cellular structure in coordinating these pathways in lung development is yet unknown. It is likely that PC act as “antennae,” sensing and transducing multiple paracrine signals that guide cell differentiation and embryonic morphogenesis. In this study, we show they are abundant throughout the lung.

We note that the markers used in this study do not definitively distinguish these embryonic structures as either primary or motile cilia. Only careful transmission electron microscopy (TEM) can do this, by assessing the structure of the (9+2) outer microtubule doublets with 2 central microtubules, something not possible with techniques used here. Generally, motile cilia are associated with expression of FoxJ1 in the cell body and DNAH11 in the cilium, among other markers (Fig. S4, S5).^24^Further detailed studies would be needed to confirm the precise types of cilia in embryonic tissues, however given those markers used here, we are likely observing the widespread presence of PC in embryonic lung tissues.

### Regionalization of epithelial ciliation and multiciliation during development

In the lung, the scientific and clinical focus has been primarily on the multiciliated cells of the upper airway that serve to sweep the airways clean of debris.^39^ When that fails, whether due to lack of motility (as in primary ciliary dyskinesia) or thickened secretions that impair flow (as in cystic fibrosis), progressive and devastating infections can result.^40^ The adult bronchial epithelium is highly heterogenous, with a variety of highly specified cell types for secreting mucus, moving the mucus along, and so forth. Here we show that the epithelial cell layer that gives rise to these upper airways initially begins as a pseudostratified but monociliated cell layer, and diverges — as physiologically necessary — along a proximal-to-distal axis.

Proximally, the main function of the epithelium is to be a barrier. Distally — at the alveolar level — its goal is to be as little a barrier as possible, providing structural support while facilitating gas exchange. The known role of FOXJ1 in driving multiciliation^41^ is observed by our immunolocalization studies at E16.5, where it expressed in a clear proximal-to-distal gradient. Epithelial cells either express Foxj1 and become multiciliated, or they do not and thus lose their PC as they further differentiate. The dynamic appearance of these different types of ciliation along the pulmonary lung epithelium is indeed interesting, and future lineage studies will be needed to define whether distinct or common cells of origin give rise to these distinct ciliated cell types. Recent descriptions of gradual deconstruction of PC in differentiated granule cells in the cerebellum point to a generality of a lack of PC in diverse tissue contexts.^42^

### Widespread absence of pulmonary endothelial cilia

ECs are highly adapted to sense and withstand the significant mechanical forces induced by blood flow (shear, stretch, pressure). PC are known to participate in force sensing and transduction,^43^ and so we expected to find cilia in pulmonary endothelial cells. Indeed, the presence of ciliated endothelial cells has been thoroughly demonstrated in the zebrafish.^44–46^ Because EC PC are observed *in vitro* under static or low shear conditions, but are either sheared off^47^ or disassembled^48^ by the onset of high shear stress flow, we predicted that cilia would be present in the capillaries and absent in the larger arteries. Surprisingly, the exact reverse was true.

Shear stress in the pulmonary arteries increases dramatically after birth. It then decreases somewhat with closure of the ductus arteriosus, and is further modulated by the continued vascular development and alveolarization that takes place in the first six months of human life. The effects of these hemodynamic changes on endothelial ciliation are not known but will be investigated in our future work. Recent computational modeling of shear stress in the pulmonary vasculature provides evidence that, in fact, shear in the distal arteries is much *higher* than in the proximal arteries, which may explain our preliminary findings on this issue.^49^ This model (higher shear stress in smaller vessels leads to loss of PC) would support a hypothesis that shear forces determine EC ciliary presence in the mouse. However, blood flow is quite minimal in early stages of development examined in our work—at which point ECs are still devoid of PC. More work is needed to understand how a primary cilium can be maintained under variable shear stresses *in vivo* — and what purpose (mechanosensitive or otherwise) the PC serve in the larger arteries.

The single cell RNA sequencing data (**Fig. S7**) reveal scattered *ARL13B* expression in ECs across time points (albeit with very little expression at E12). This expression could indicate either i) cilia formation that we failed to see by IF, ii) *Arl13b* transcription and localization not associated with cilia,^50^ iii) contamination by ambient RNA, or iv) imperfect separation of closely associated endothelial and mesenchymal cells during processing for scRNAseq, as previously reported in brain parenchyma.^51^ Indeed, prior work has claimed the identification of primary cilia in a single human pulmonary arterial ECs (Fig 1H from Lee 2018).^52^ While it is difficult to tell, this EC appears to be in a larger artery. While we are confident in our analysis of the IF stains, further work is needed to clarify the expression and role of ciliary genes and proteins in ECs in the larger arteries of the adult human lung.

### Mesenchymal cilia: What is their function?

The ubiquity of PC in the mesenchyme of the developing lung that we found is remarkable, and highlights yet again the necessity of *in vivo* studies. *In vitro*, PC are typically only found on cultured cells that have been serum-starved in order to enter a quiescent state. Pulmonary mesenchymal cells by contrast are actively growing, migrating, and dividing during active organogenesis, and are thus anything but static or quiescent. Future studies will be needed to investigate the presence of PC throughout the cell cycle, and the role cilia play in cellular migration or differentiation during organogenesis.

### Single cell analysis reveals lung mesenchyme heterogeneity and identity of ciliated cells

Understanding exactly what cell type builds and maintains PC in the lung is likely to be critical for a mechanistic understanding of ciliopathy phenotypes of the lung. As the current study demonstrates, it is no small feat identifying with clarity the spatiotemporal cell-of-origin of the cilia. At this time, there is no clear gene signature for cells with PC that could be used to track their status using single cell sequencing methods. We have chosen to examine ARL13Bas a proxy for ciliated cells. However, transcript counts of *Arl13b* are not necessarily true hallmarks of a ciliated cell — as this measure likely reflects cells in the process of active cilium generation. However, our immunofluorescent localization pointed to the lung stroma as a likely population of cells harboring PC. This, in parallel with our single cell analysis of the lung mesenchyme, allowed us to begin painting a picture of which cell populations construct and maintain PC over developmental time. Lack of maintenance of ciliary and peri-ciliary factors arising from transcriptional regulation has also been recently reported in differentiated granule cells of the cerebellum that lack cilia.^53^

It is important to note that postnatal development continues in many organs — and it is *critical* for the lung. After birth, alveologenesis continues up to P36 in mice to create the mature, functional lung. Secondary crest myofibroblasts (SCMFs) play a critical role in alveologenesis by providing the contractile force to generate the “secondary crest” that grows into a new, mature alveolar wall, as well as providing crucial paracrine signals during septae formation.^54,55^ Building on the previously reported importance of PDGFRα signaling to SCMFs, we now find that SCMFs widely and robustly exhibit PC. We are currently investigating the role of various ciliary signaling pathways in this population. Additionally, the fact that a remarkably active, contractile cell population like the SCMFs all display PC suggests a role in directed cellular migration and cell-cell signaling. We might soon come to recognize the PC not just as an antenna, but also a rudder, steering the cellular ship and keeping it on course.

## Acknowledgements

We gratefully acknowledge the UT Southwestern Electron Microscopy Core Facility (Natalia Gunko, Ph.D. Equipment purchased by 1S10OD021685-01A1 to Katherine Luby-Phelps), for tissue processing and imaging training/assistance with TEM. We thank the UT Southwestern Quantitative Light Microscopy Core Facility (Marcel Mettlen, Ph.D.), for training/assistance. Additional TEM imaging work was performed at the Northwestern University Center for Advanced Microscopy (RRID: SCR_020996) generously supported by NCI CCSG P30 CA060553 awarded to the Robert H Lurie Comprehensive Cancer Center. We are also grateful to Michael Dellinger, Ph.D. for generous supply of reagents.

## Author contributions

Conceptualization: S.S., O.C.; Methodology: S.S., O.C.; Formal analysis: S.S., M.N.; Investigation: S.S., M.N., J.S., F.N.C., S.N.M.; Resources: O.C..; Data curation: S.S., M.N., F.N.C., S.N.M.; Writing - original draft: S.S., O.C.; Writing - review & editing: S.S., M.N., F.N.C., S.N.M., M.L.I.A., J.A.Z., S.M., O.C.; Visualization: S.S., M.N.; Supervision: S.M., O.C.; Project administration: S.S.; Funding acquisition: O.C.

## Supplementary Figure Legends

**Fig. S1.**
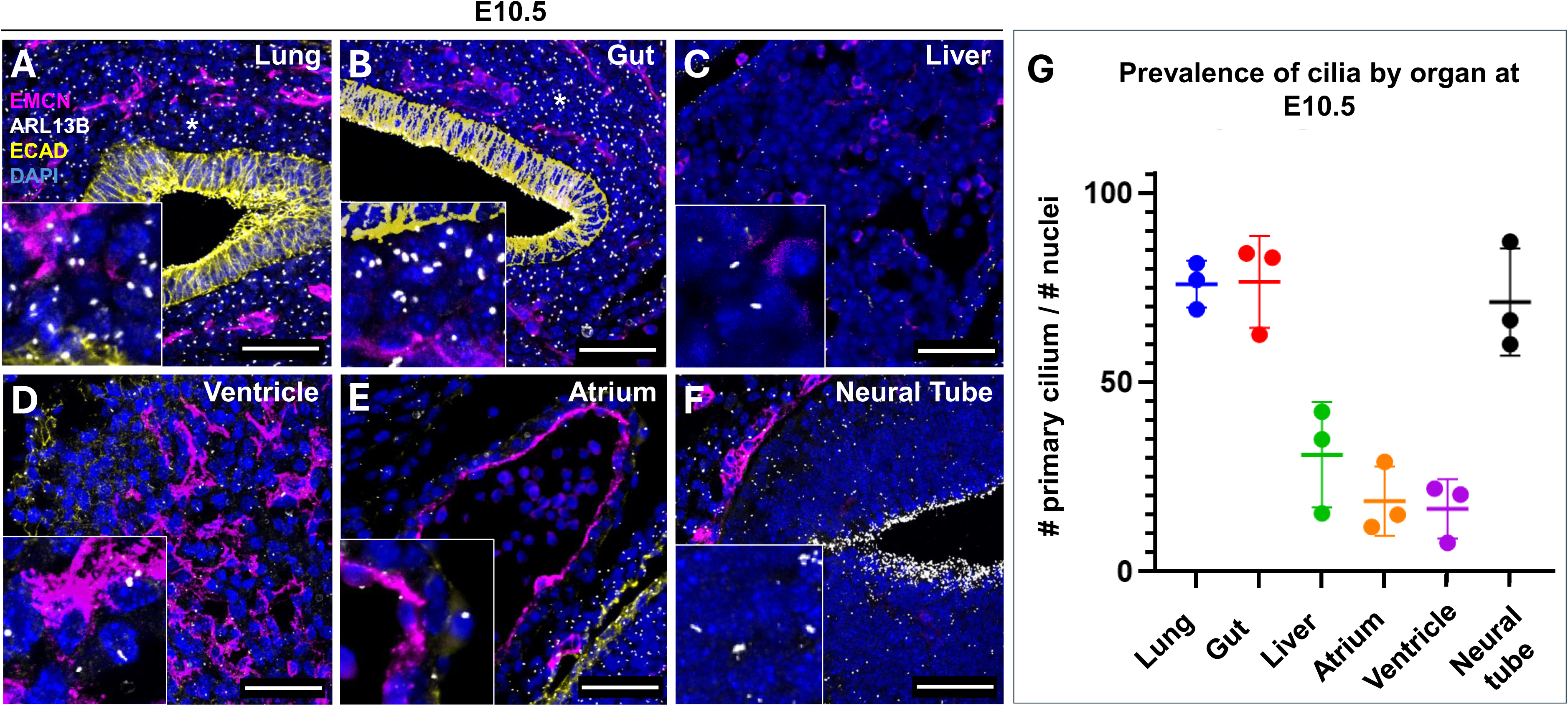
Widespread, multi-organ presence of primary cilia during organogenesis. Immunofluorescent staining of 10 µm slices of paraffin-fixed, whole E10.5 embryos (n=3) were used for whole-organ assessments of overall ciliary density in the all cells of developing lung **(A)**, gut **(B)**, liver **(C)**, cardiac ventricle **(D)**, atrium **(E)**, and neural tube **(F)**. Total number of cilia (ARL13B, white) relative to the total number of nuclei (DAPI) was quantified in 3x digitally magnified images of 20x scans **(G)**. Scale bar is 50 µm.

**Fig. S2.**
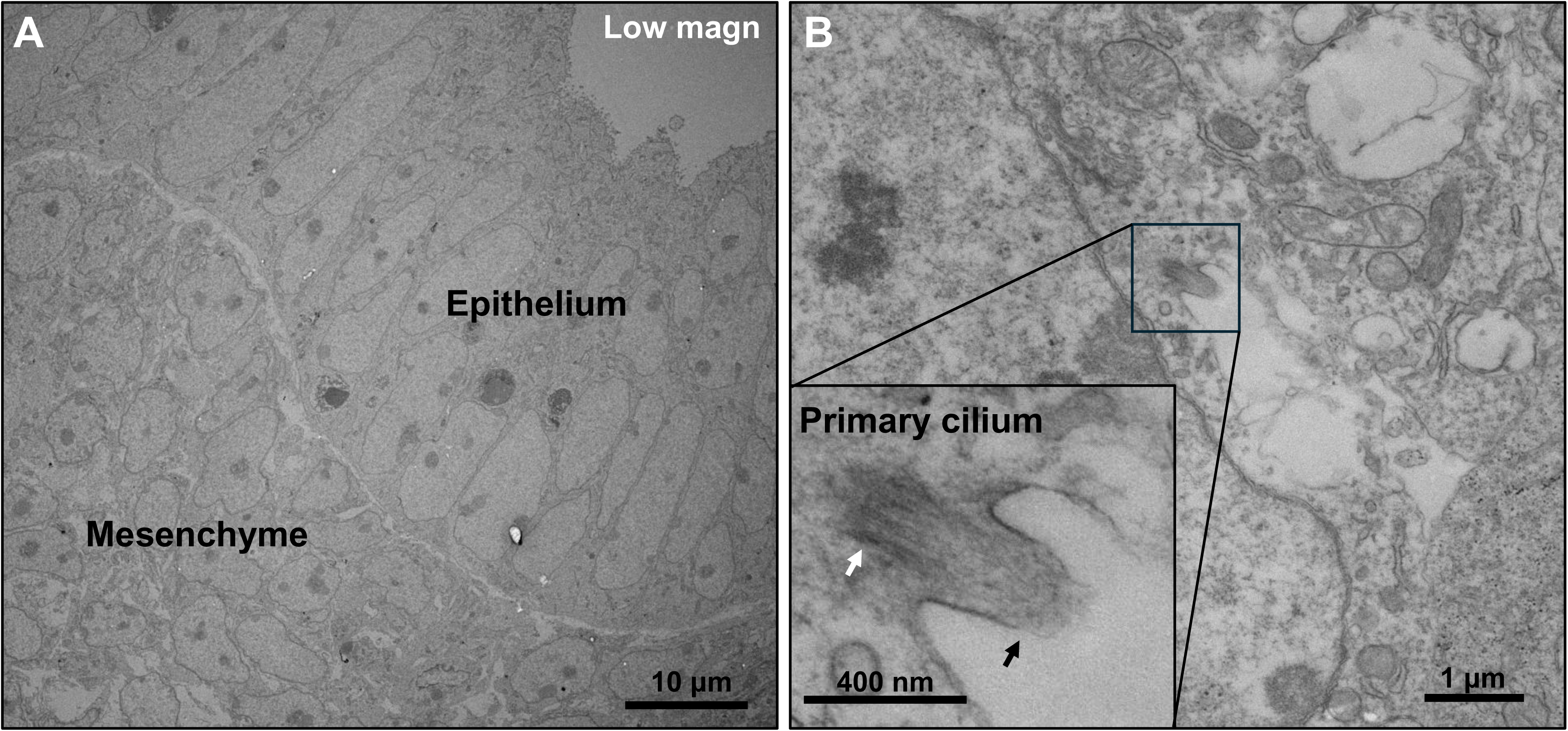
Confirmation of mesenchymal primary cilia in the lung by electron microscopy. Whole E12.5 embryos were fixed in 4% PFA for TEM. **(A)** Interface of pseudostratified pulmonary epithelium and mesenchyme shown. **(B)** High magnification view of a primary cilium in the lung mesenchyme. Inset shows detail of a primary cilium (white arrow = basal body, black arrow = ciliary axoneme, sectioned). The cilium sits within a deep ciliary pocket. Scale bars sizes as indicated.

**Fig. S3.**
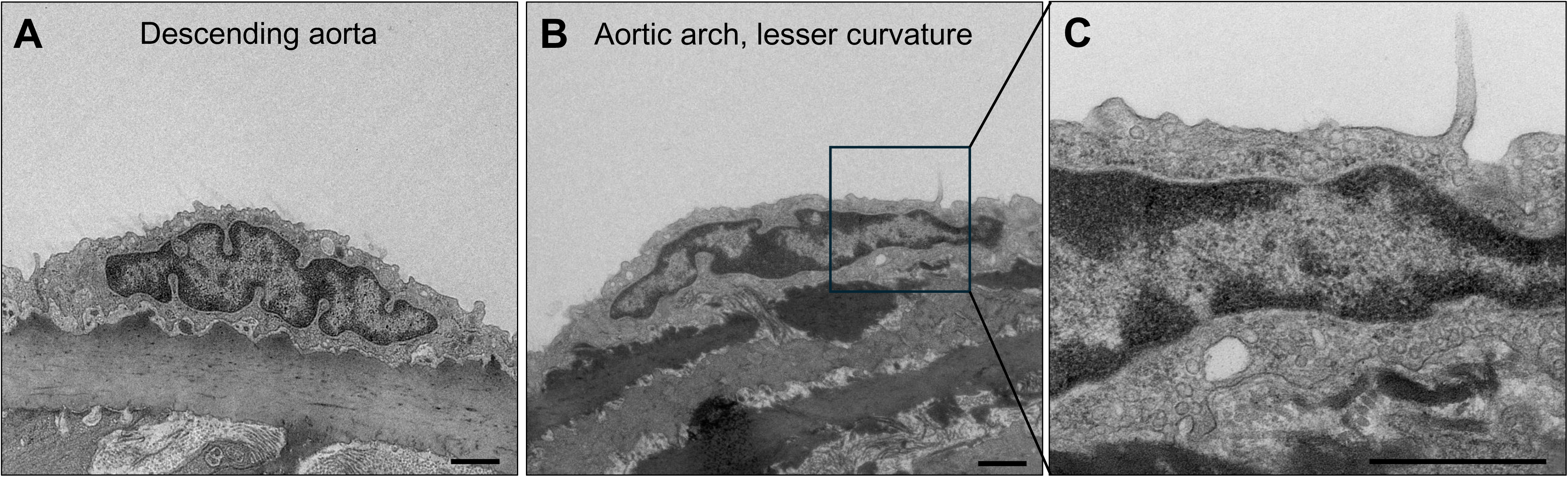
Confirmation of endothelial primary cilia in the aorta by electron microscopy. **(A)** Adult aortic ECs in the region of the descending aorta (P60) were not found to express PC. The lesser curvature of the aortic arch contains some ECs with primary cilia, shown at **(B)** low magnification and **(C)** high magnification. Scale bar is 1 μm.

**Fig. S4.**
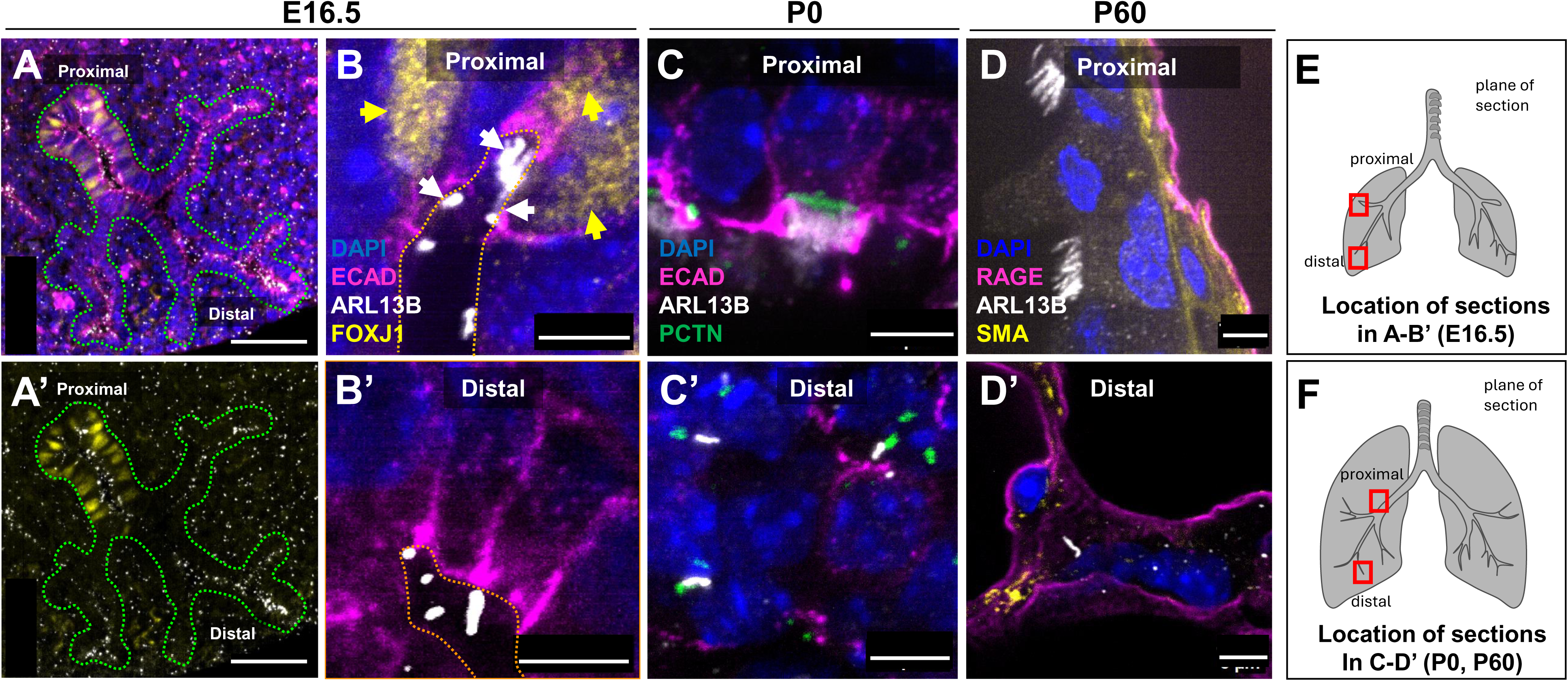
Proximal-distal gradient of ciliary fate in the developing pulmonary epithelium. Immunofluorescent analysis of sectioned lung tissue from embryonic **(A-B’)**, P0 **(C,C’)** and P60 **(D,D’)**. **(A-B’)** Note higher levels of FoxJ1 in proximal epithelium than in distal epithelium, and FoxJ1+ cells exhibit different stages of multiciliation **(B).** Proximal and distal regions as indicated. By contrast, cells lacking FoxJ1 display single PC or no PC, particularly in the middle portions of the airway **(A’, B’)**. Similarly, ciliated cells of proximal epithelium exhibit multicilia at P0 **(C)** and P60 **(D)**, while distal epithelial cells (positive for either ECAD or RAGE) exhibit single primary cilia **(C’, D’)**. Epithelial basement membrane is outlined by green dotted line in **(A),** and alveolar lumen is outlined by orange dotted line in **(B)**. Scale bar is 5 μm except in **A, A’** where bar is 50 μm.

**Fig. S5.**
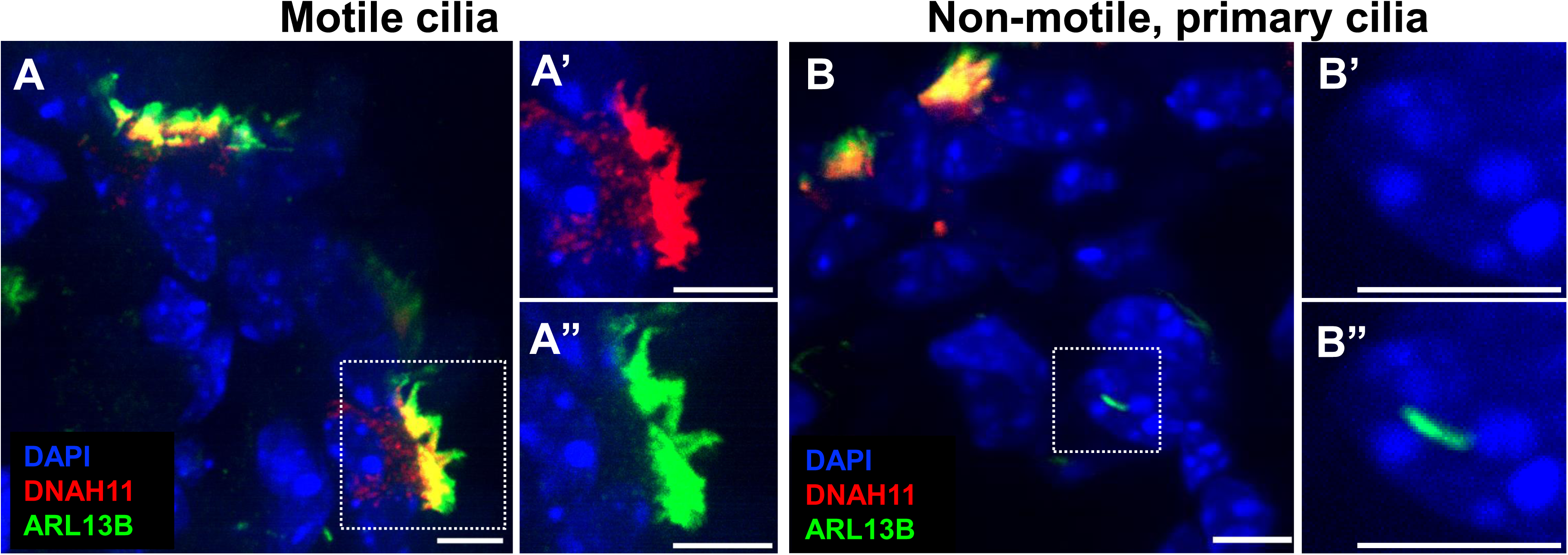
Expression of DNAH11 specifically within motile cilia. Immunofluorescent staining of P30 mouse tissue for DNAH11, a key component of motile cilia that is not present in primary cilia, shows clearly that the mesenchymal cilia identified in this study do not have DNAH11 and are likely non-motile. **(A)** Motile cluster of cilia showing co-expression of DNAH11 and ARL13B in upper airway epithelial cells. **(B)** Primary cilia in alveolar epithelium (more distal) showing lack of DNAH11 expression. Note nearby multiciliated cell at top of panel in B. Insets (dotted white line) show both red and green channels in **A’, A”, B’, B”**, showing details of multiciliated and primary ciliary structures, as well as presence or absence of staining. Scale bar is 5 μm.

**Figure S6.**
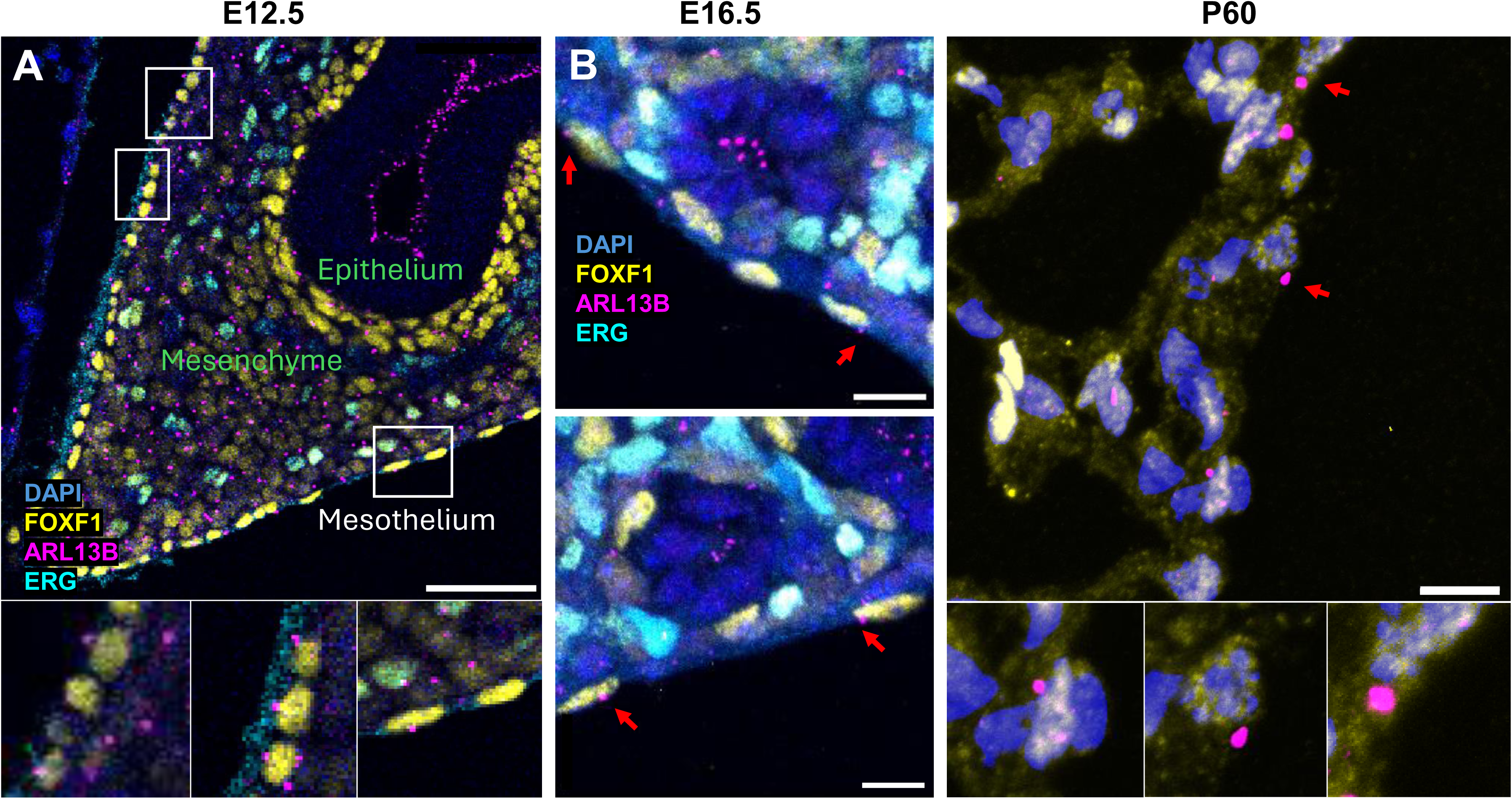
Mesothelial cells lining the periphery of the lung harbor PC. Immunofluorescent analysis of lung mesothelium at various stages of development revealed ciliated mesothelial cells. Not all mesothelial cells appear ciliated in these 10 μm slices. FOXF1 is expressed in ciliated mesothelial cells at E12.5 **(A)**, E16.5 **(B)**, and P60 **(C).** Red arrows highlight primary cilia (ARL13B, magenta)

**Fig. S7.**
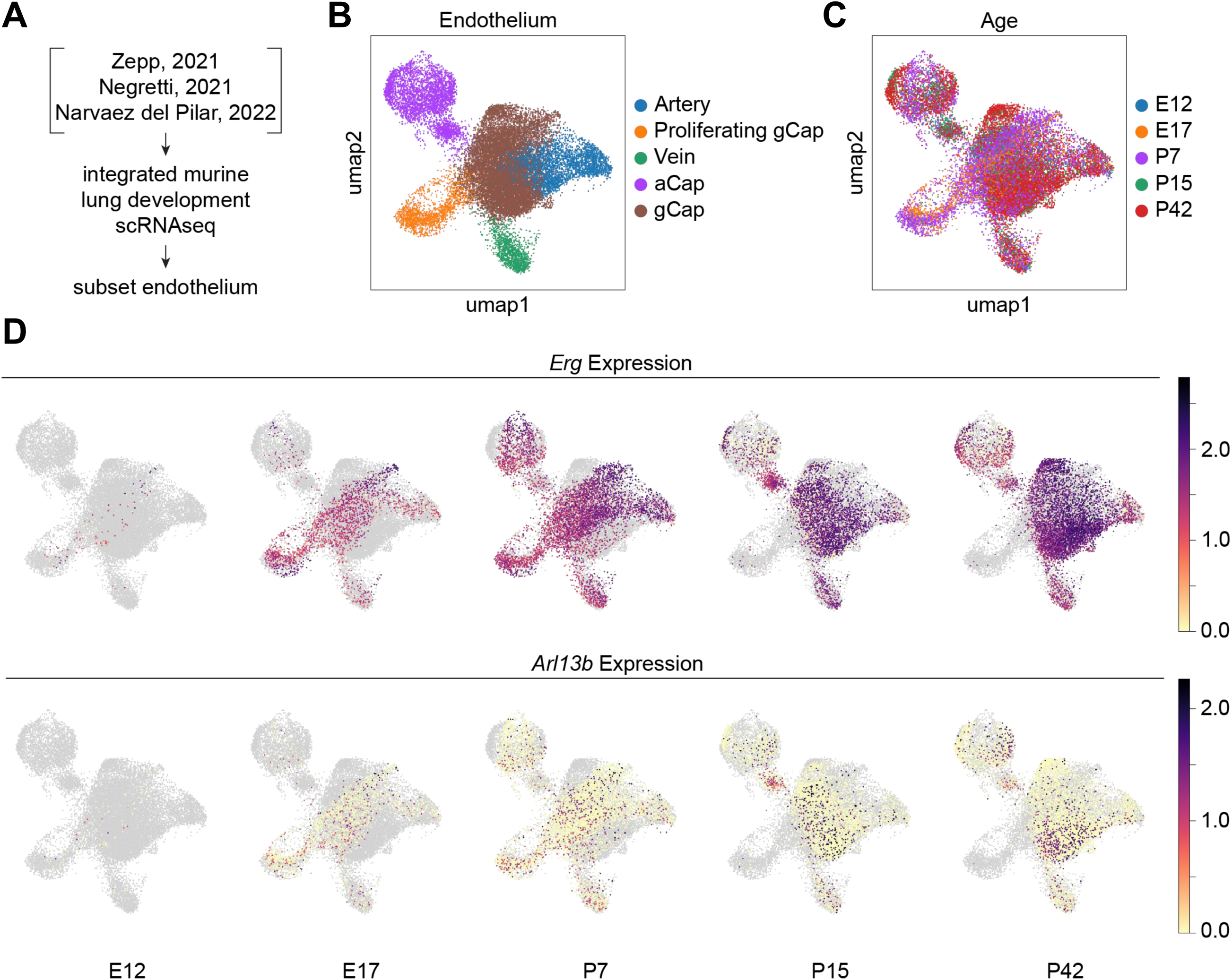
Capillary *Arl13b* expression found in the developing endothelium. **(A)** Previously published lung development scRNAseq datasets were integrated and the endothelium was subset. **(B)** UMAP of endothelial cell types **(C)** UMAP of developmental timepoints. **(D)** Feature plots show expression of *Erg* and *Arl13b* in the lung endothelium across developmental timepoints.

**Fig. S8.**
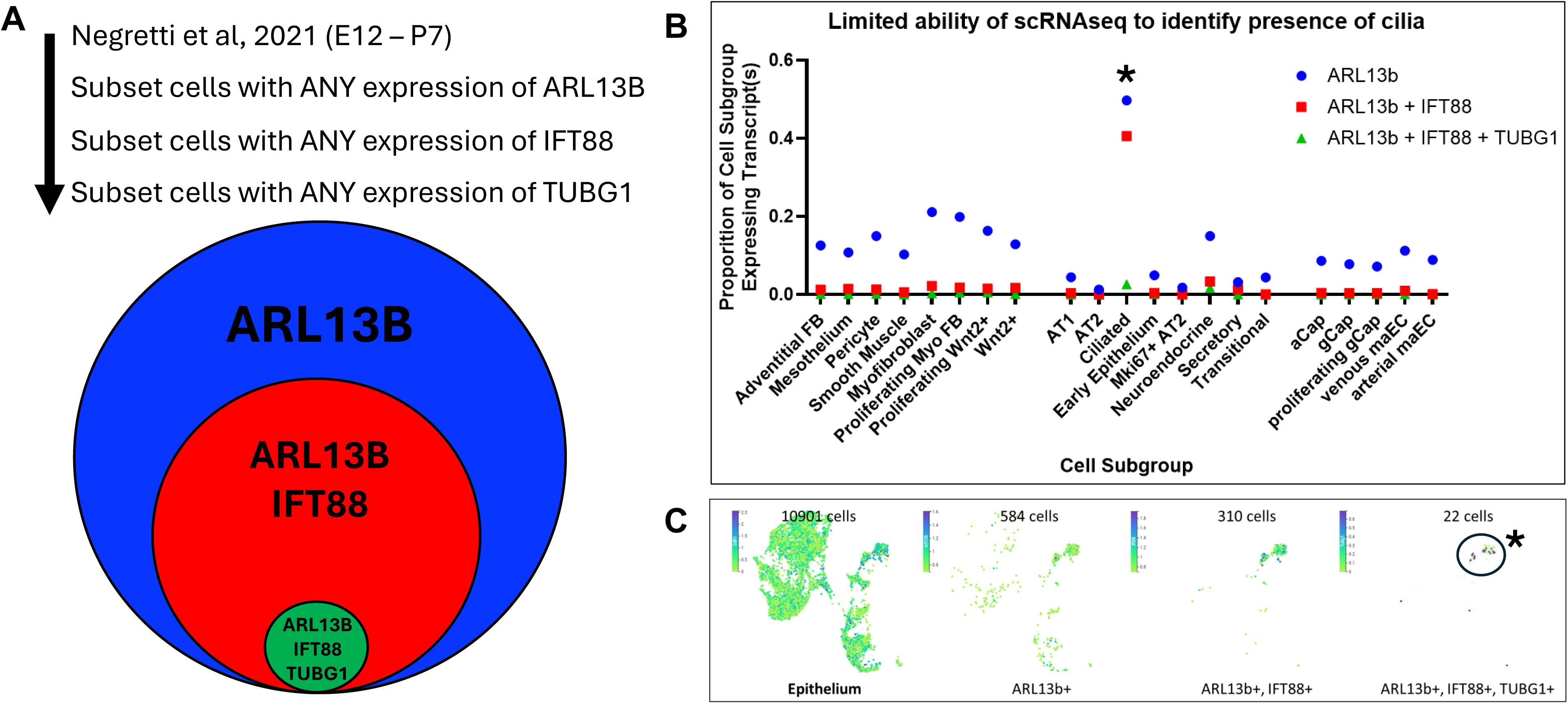
Limited ability of gene sets to identify ciliated cells in single cell RNA sequencing. **(A)** By sequentially identifying cells with any expression of *Arl13b*, *Ift88*, and *Tugb1*, we assessed whether single cell data could be used for subsequent pathway analysis of ciliated cells. **(B)** Cells identified as ciliated (**\***) by overall pathway analysis (Negretti 2021) only showed active transcription of *Arl13b* in 50% of cells, while only 41% of cells were double positive for *Arl13b* and *Ift88*, and 2% were positive for all three (triple positive cells, circled and indicated with asterisk. **(C)** Given clear co-localization in cilia in our immunofluorescent analysis, scRNAseq data was not used for subsequent analysis of known gene ontology and pathway analysis.

**Fig. S9.**
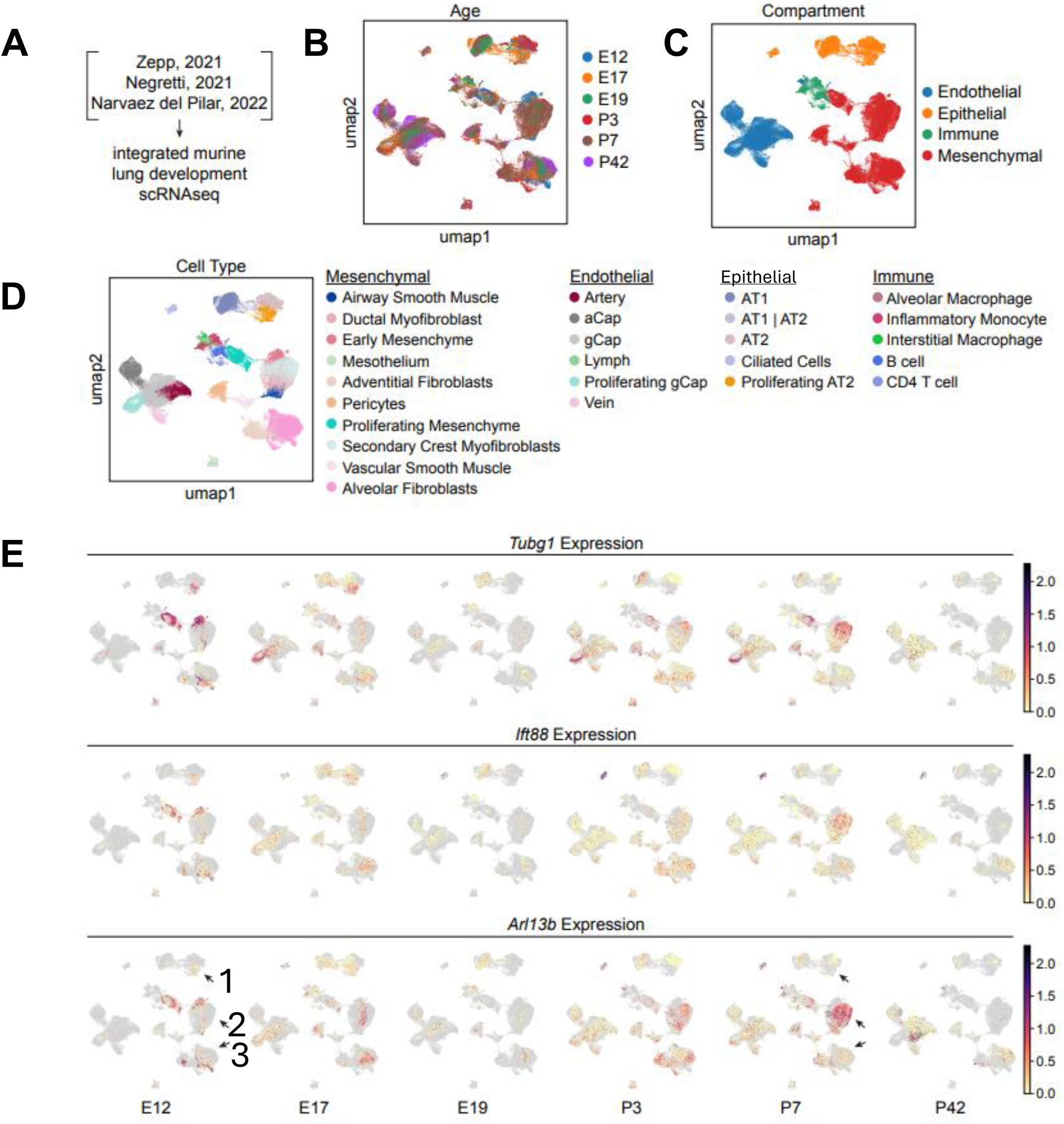
Single cell mRNA sequencing is useful to highlight active ciliogenesis but not static ciliary presence. **(A)** Previously published lung development scRNA seq datasets were integrated. **(B)** UMAP of developmental time points **(C)** UMAP of major cell groups. **(D)** UMAP of detailed cellular subgroups. **(E)** Feature plots show expression of *Tubg1, Ift88,* and *Arl13b* in the lung mesenchyme across developmental timepoints. Arrows highlight *Arl13b* expression in ciliated epithelial cells (1), secondary crest myofibroblasts (2), and alveolar fibroblasts (3).

**Table S1.**

| **Target Protein** | **Host Species** | **Manufacturer** | **Catalog No.** | **Dilution Factor** |
| --- | --- | --- | --- | --- |
| Alpha Tubulin | Rabbit | Cell Signaling Technology | 5335T | 1 to 100 |
| ARL13B | Rabbit | Protein Tech | 17711-1-AP | 1 to 200 |
| CA4 | Goat | R&D Systems | AF2414 | 1 to 100 |
| DNAH11 | Mouse | Thermofisher | MA5-43853 | 1 to 100 |
| E-cadherin | Rat | Invitrogen | 13-1900 | 1 to 100 |
| Endomucin | Rat | Santa Cruz | sc-65495 | 1 to 100 |
| ERG 1/2/3 | Mouse | Santa Cruz | sc-376293 | 1 to 100 |
| FOXJ1 | Mouse | Invitrogen | 14-9965-82 | 1 to 100 |
| Gamma Tubulin | Rabbit | Abcam | ab11317 | 1 to 100 |
| GFP | Chicken | Aves Labs | GFP-1020 | 1 to 500 |
| GM130 | Mouse | BD Biosci | 610823 | 1 to 200 |
| GM130 | Rat | BioLegend | 937001 | 1 to 100 |
| HOP | Mouse | Santa Cruz | sc-398703 | 1 to 50 |
| IFT88 | Rabbit | Proteintech | 13967-1-AP | 1 to 100 |
| LAMP3 | Rat | Proteintech | 12632-1-AP | 1 to 100 |
| PDGFRα | Goat | R&D Systems | AF1062 | 1 to 100 |
| PDGFRβ | Rat | Invitrogen | 14-1402-82 | 1 to 100 |
| Pericentrin | Mouse | BD Biosci | 611814 | 1 to 100 |
| PROX1 | Goat | R&D Systems | AF2727 | 1 to 100 |
| RAGE | Rat | R&D Systems | MAB1179 | 1 to 100 |
| SMA Cy3 | Mouse | Sigma | C6198 | 1 to 400 |
| Sox17 | Goat | R&D Systems | AF1924 | 1 to 100 |
| Ve-cadherin | Mouse | Santa Cruz | sc-9989 | 1 to 100 |

**Table S2.**
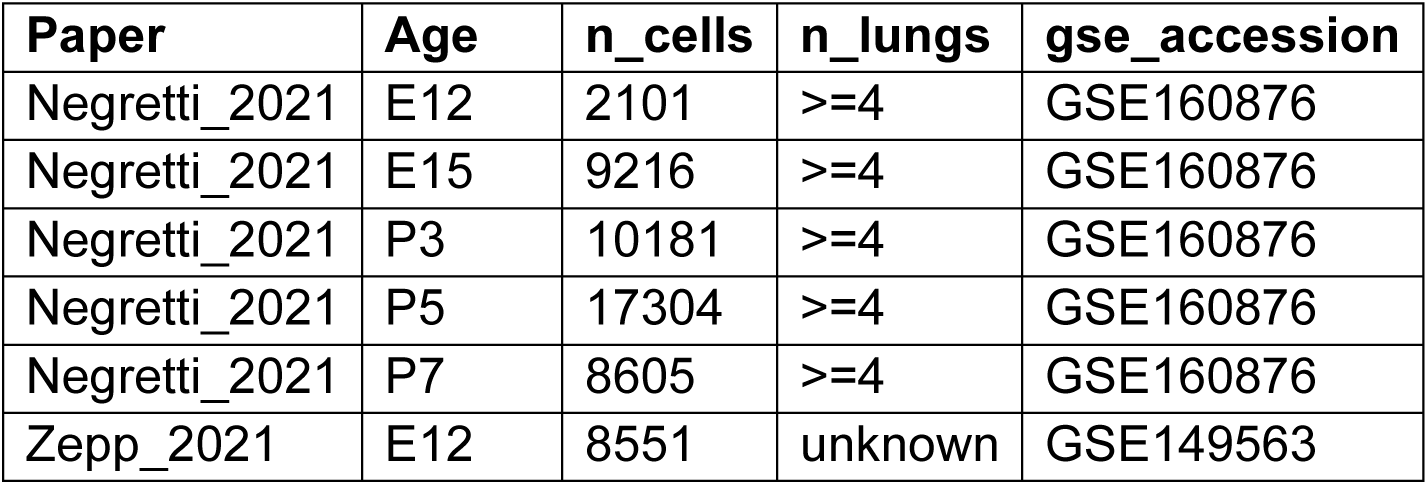
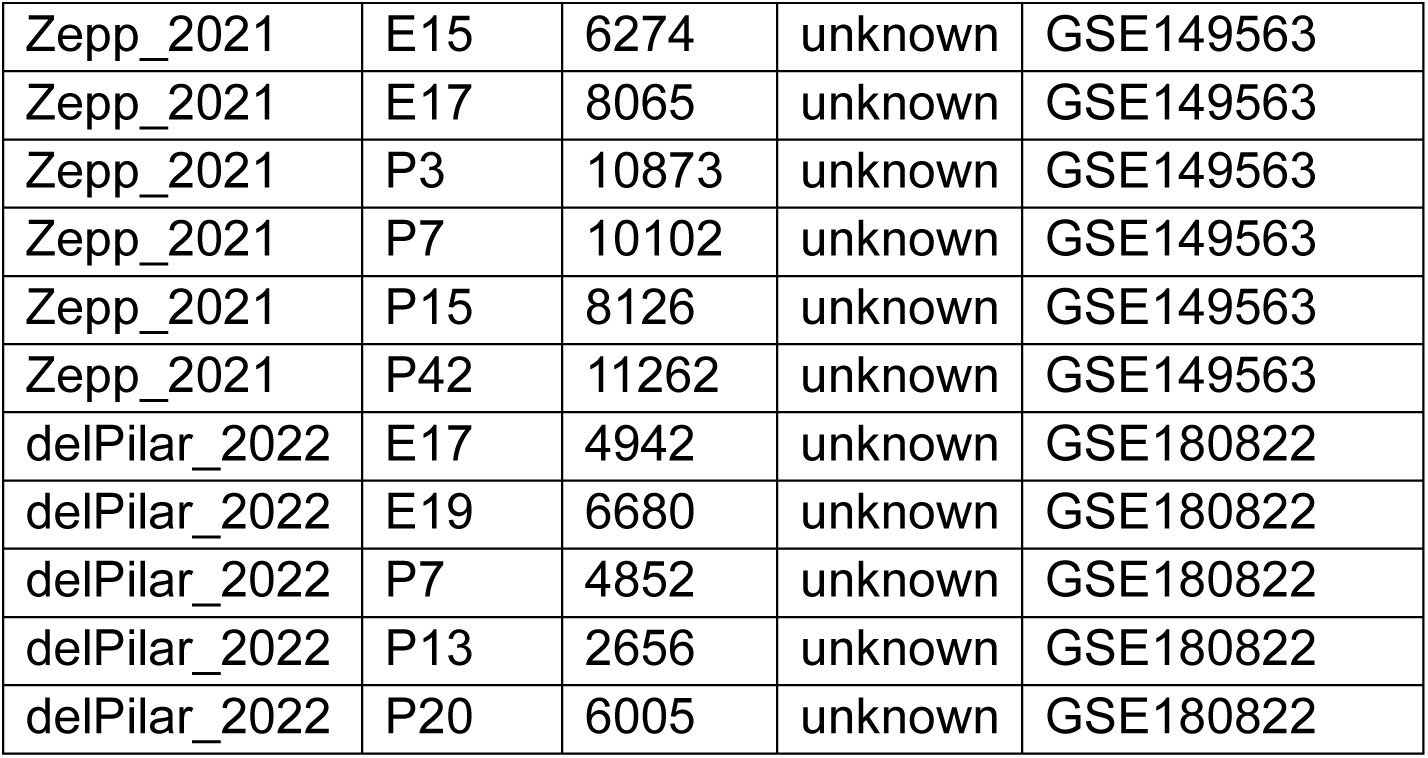

## References

1. Kim S, Dynlacht BD. Assembling a primary cilium. Curr Opin Cell Biol. Aug 2013;25(4):506–11. 10.1016/j.ceb.2013.04.011.

2. Nonaka S, Shiratori H, Saijoh Y, Hamada H. Determination of left-right patterning of the mouse embryo by artificial nodal flow. Nature. Jul 4 2002;418(6893):96-9. 10.1038/nature00849.

3. Clement CA, Kristensen SG, Mollgard K, et al. The primary cilium coordinates early cardiogenesis and hedgehog signaling in cardiomyocyte differentiation. J Cell Sci. Sep 1 2009;122(Pt 17):3070–82. 10.1242/jcs.049676.

4. Nonaka S, Tanaka Y, Okada Y, et al. Randomization of left-right asymmetry due to loss of nodal cilia generating leftward flow of extraembryonic fluid in mice lacking KIF3B motor protein. Cell. Dec 11 1998;95(6):829–37. 10.1016/s0092-8674(00)81705-5.

5. Li S, Zhang H, Sun Y. Primary cilia in hard tissue development and diseases. Front Med. Oct 2021;15(5):657–678. 10.1007/s11684-021-0829-6.

6. Haycraft CJ, Zhang Q, Song B, et al. Intraflagellar transport is essential for endochondral bone formation. Development. Jan 2007;134(2):307–16. 10.1242/dev.02732.

7. Bakey Z, Cabrera OA, Hoefele J, et al. IFT74 variants cause skeletal ciliopathy and motile cilia defects in mice and humans. PLoS Genet. Jun 2023;19(6):e1010796. 10.1371/journal.pgen.1010796.

8. Klena N, Gabriel G, Liu X, et al. Role of Cilia and Left-Right Patterning in Congenital Heart Disease. In: Nakanishi T, Markwald RR, Baldwin HS, Keller BB, Srivastava D, Yamagishi H, eds. Etiology and Morphogenesis of Congenital Heart Disease: From Gene Function and Cellular Interaction to Morphology. Tokyo: 2016:67–79.

9. Willaredt MA, Hasenpusch-Theil K, Gardner HA, et al. A crucial role for primary cilia in cortical morphogenesis. J Neurosci. Nov 26 2008;28(48):12887–900. 10.1523/JNEUROSCI.2084-08.2008.

10. Huangfu D, Anderson KV. Cilia and Hedgehog responsiveness in the mouse. Proc Natl Acad Sci U S A. Aug 9 2005;102(32):11325–30. 10.1073/pnas.0505328102.

11. Amack JD. Structures and functions of cilia during vertebrate embryo development. Mol Reprod Dev. Dec 2022;89(12):579–596. 10.1002/mrd.23650.

12. Pal K, Mukhopadhyay S. Primary cilium and sonic hedgehog signaling during neural tube patterning: role of GPCRs and second messengers. Dev Neurobiol. Apr 2015;75(4):337–48. 10.1002/dneu.22193.

13. Cano DA, Sekine S, Hebrok M. Primary Cilia Deletion in Pancreatic Epithelial Cells Results in Cyst Formation and Pancreatitis. Gastroenterology. 2006;131(6):1856–1869. 10.1053/j.gastro.2006.10.050.

14. Jain R, Pan J, Driscoll JA, et al. Temporal Relationship between Primary and Motile Ciliogenesis in Airway Epithelial Cells. American Journal of Respiratory Cell and Molecular Biology. 2010;43(6):731–739. 10.1165/rcmb.2009-0328oc.

15. Azizoglu DB, Braitsch C, Marciano DK, Cleaver O. Afadin and RhoA control pancreatic endocrine mass via lumen morphogenesis. Genes & Development. 2017;31(23-24):2376–2390. 10.1101/gad.307637.117.

16. Chaudhry FN, Michki NS, Shirmer DL, et al. Dynamic Hippo pathway activity underlies mesenchymal differentiation during lung alveolar morphogenesis. Development. Apr 15 2024;151(8). 10.1242/dev.202430.

17. Narvaez Del Pilar O, Gacha Garay MJ, Chen J. Three-axis classification of mouse lung mesenchymal cells reveals two populations of myofibroblasts. Development. Mar 15 2022;149(6). 10.1242/dev.200081.

18. Zepp JA, Morley MP, Loebel C, et al. Genomic, epigenomic, and biophysical cues controlling the emergence of the lung alveolus. Science. Mar 12 2021;371(6534). 10.1126/science.abc3172.

19. Negretti NM, Plosa EJ, Benjamin JT, et al. A single-cell atlas of mouse lung development. Development. Dec 15 2021;148(24). 10.1242/dev.199512.

20. Kaminow B, Yunusov D, Dobin A. STARsolo: accurate, fast and versatile mapping/quantification of single-cell and single-nucleus RNA-seq data [Preprint]. BioRXiv. May 5 2021. 10.1101/2021.05.05.442755.

21. Virshup I, Bredikhin D, Heumos L, et al. The scverse project provides a computational ecosystem for single-cell omics data analysis. Nat Biotechnol. May 2023;41(5):604–606. 10.1038/s41587-023-01733-8.

22. Kasahara K, Miyoshi K, Murakami S, Miyazaki I, Asanuma M. Visualization of astrocytic primary cilia in the mouse brain by immunofluorescent analysis using the cilia marker Arl13b. Acta Med Okayama. Dec 2014;68(6):317–22. 10.18926/AMO/53020.

23. Avasthi P, Marshall WF. Stages of ciliogenesis and regulation of ciliary length. Differentiation. 2012;83(2):S30–S42. 10.1016/j.diff.2011.11.015.

24. Shoemark A, Frost E, Dixon M, et al. Accuracy of Immunofluorescence in the Diagnosis of Primary Ciliary Dyskinesia. American Journal of Respiratory and Critical Care Medicine. 2017;196(1):94–101. 10.1164/rccm.201607-1351oc.

25. Klena N, Pigino G. Structural Biology of Cilia and Intraflagellar Transport. Annu Rev Cell Dev Biol. Oct 6 2022;38:103–123. 10.1146/annurev-cellbio-120219-034238.

26. Graeme, Aarti, Dufton N, et al. The Endothelial Transcription Factor ERG Promotes Vascular Stability and Growth through Wnt/β-Catenin Signaling. Developmental Cell. 2015;32(1):82–96. 10.1016/j.devcel.2014.11.016.

27. Caporarello N, Lee J, Pham TX, et al. Dysfunctional ERG signaling drives pulmonary vascular aging and persistent fibrosis. Nat Commun. Jul 25 2022;13(1):4170. 10.1038/s41467-022-31890-4.

28. Gillich A, Zhang F, Farmer CG, et al. Capillary cell-type specialization in the alveolus. Nature. Oct 2020;586(7831):785–789. 10.1038/s41586-020-2822-7.

29. Plotnikova OV, Pugacheva EN, Golemis EA. Primary Cilia and the Cell Cycle. Elsevier; 2009:137–160.

30. Dinsmore C, Reiter JF. Endothelial primary cilia inhibit atherosclerosis. EMBO Rep. Feb 2016;17(2):156–66. 10.15252/embr.201541019.

31. Trempus CS, Song W, Lazrak A, et al. A novel role for primary cilia in airway remodeling. Am J Physiol Lung Cell Mol Physiol. Aug 1 2017;313(2):L328–L338. 10.1152/ajplung.00284.2016.

32. Weibel ER. Morphological basis of alveolar-capillary gas exchange. Physiol Rev. Apr 1973;53(2):419–95. 10.1152/physrev.1973.53.2.419.

33. Mason RJ, Williams MC. Type II alveolar cell. Defender of the alveolus. Am Rev Respir Dis. Jun 1977;115(6 Pt 2):81–91. 10.1164/arrd.1977.115.S.81.

34. Davis JD, Wypych TP. Cellular and functional heterogeneity of the airway epithelium. Mucosal Immunol. Sep 2021;14(5):978–990. 10.1038/s41385-020-00370-7.

35. El Agha E, Thannickal VJ. The lung mesenchyme in development, regeneration, and fibrosis. J Clin Invest. Jul 17 2023;133(14). 10.1172/JCI170498.

36. Morrisey EE, Hogan BL. Preparing for the first breath: genetic and cellular mechanisms in lung development. Dev Cell. Jan 19 2010;18(1):8–23. 10.1016/j.devcel.2009.12.010.

37. Caprioli A, Villasenor A, Wylie LA, et al. Wnt4 is essential to normal mammalian lung development. Dev Biol. Oct 15 2015;406(2):222–34. 10.1016/j.ydbio.2015.08.017.

38. Goetz SC, Anderson KV. The primary cilium: a signalling centre during vertebrate development. Nat Rev Genet. May 2010;11(5):331–44. 10.1038/nrg2774.

39. Rawlins EL, Ostrowski LE, Randell SH, Hogan BL. Lung development and repair: contribution of the ciliated lineage. Proc Natl Acad Sci U S A. Jan 9 2007;104(2):410–7. 10.1073/pnas.0610770104.

40. Tilley AE, Walters MS, Shaykhiev R, Crystal RG. Cilia dysfunction in lung disease. Annu Rev Physiol. 2015;77:379–406. 10.1146/annurev-physiol-021014-071931.

41. Blatt EN, Yan XH, Wuerffel MK, Hamilos DL, Brody SL. Forkhead transcription factor HFH-4 expression is temporally related to ciliogenesis. Am J Respir Cell Mol Biol. Aug 1999;21(2):168–76. 10.1165/ajrcmb.21.2.3691.

42. Ott CM, Constable S, Nguyen TM, et al. Permanent deconstruction of intracellular primary cilia in differentiating granule cell neurons. J Cell Biol. Oct 7 2024;223(10). 10.1083/jcb.202404038.

43. Satir P, Christensen ST. Overview of structure and function of mammalian cilia. Annu Rev Physiol. 2007;69:377–400. 10.1146/annurev.physiol.69.040705.141236.

44. Jacky, Steed E, Rita, et al. Endothelial Cilia Mediate Low Flow Sensing during Zebrafish Vascular Development. Cell Reports. 2014;6(5):799–808. 10.1016/j.celrep.2014.01.032.

45. Supatto W, Vermot J. From cilia hydrodynamics to zebrafish embryonic development. Curr Top Dev Biol. 2011;95:33–66. 10.1016/B978-0-12-385065-2.00002-5.

46. Eisa-Beygi S, Benslimane FM, El-Rass S, et al. Characterization of Endothelial Cilia Distribution During Cerebral-Vascular Development in Zebrafish ( *Danio rerio* ). Arteriosclerosis, Thrombosis, and Vascular Biology. 2018;38(12):2806–2818. 10.1161/atvbaha.118.311231.

47. Van der Heiden K, Hierck BP, Krams R, et al. Endothelial primary cilia in areas of disturbed flow are at the base of atherosclerosis. Atherosclerosis. Feb 2008;196(2):542–50. 10.1016/j.atherosclerosis.2007.05.030.

48. Iomini C, Tejada K, Mo W, Vaananen H, Piperno G. Primary cilia of human endothelial cells disassemble under laminar shear stress. J Cell Biol. Mar 15 2004;164(6):811–7. 10.1083/jcb.200312133.

49. Yang W, Dong M, Rabinovitch M, Chan FP, Marsden AL, Feinstein JA. Evolution of hemodynamic forces in the pulmonary tree with progressively worsening pulmonary arterial hypertension in pediatric patients. Biomechanics and Modeling in Mechanobiology. 2019;18(3):779–796. 10.1007/s10237-018-01114-0.

50. Ferent J, Constable S, Gigante ED, et al. The Ciliary Protein Arl13b Functions Outside of the Primary Cilium in Shh-Mediated Axon Guidance. Cell Reports. 2019;29(11):3356–3366.e3. 10.1016/j.celrep.2019.11.015.

51. Heumos L, Schaar AC, Lance C, et al. Best practices for single-cell analysis across modalities. Nature Reviews Genetics. 2023;24(8):550–572. 10.1038/s41576-023-00586-w.

52. Lee J, Oh DH, Park KC, et al. Increased Primary Cilia in Idiopathic Pulmonary Fibrosis. Mol Cells. Mar 31 2018;41(3):224–233. 10.14348/molcells.2018.2307.

53. Constable S, Ott CM, Lemire AL, et al. Programmed withdrawal of cilia maintenance followed by centriole capping leads to permanent cilia loss during cerebellar granule cell neurogenesis. Cold Spring Harbor Laboratory; 2023.

54. Branchfield K, Li R, Lungova V, Verheyden JM, McCulley D, Sun X. A three-dimensional study of alveologenesis in mouse lung. Dev Biol. Jan 15 2016;409(2):429–41. 10.1016/j.ydbio.2015.11.017.

55. Li R, Li X, Hagood J, Zhu M-S, Sun X. Myofibroblast contraction is essential for generating and regenerating the gas-exchange surface. Journal of Clinical Investigation. 2020;130(6):2859–2871. 10.1172/jci132189.

